# A self-complementary recombinant adeno-associated virus vector coding for an anchorless prion protein carrying the G127V mutation extends survival in a rodent prion disease model

**DOI:** 10.64898/2026.03.27.714700

**Authors:** Thomas Zerbes, Claire Verkuyl, Cunjie Zhang, Sophie Grunnesjoe, Shehab Eid, Hamza Arshad, Wenda Zhao, Zahra Nasser, Teaghan O’Shea, Ari Belotserkovsky, Lise Lamoureux, Kathy L Frost, Jennifer Myskiw, Leyao Li, Erica Stuart, Holger Wille, Stephanie Booth, Joel C. Watts, Gerold Schmitt-Ulms

## Abstract

The replacement of a single codon in the human prion gene, causing the substitution of glycine with valine at position 127 (G127V) of the prion protein (PrP), prevents development of prion disease. We set out to explore if prion disease survival extension manifests in mice if the V127 mutant is delivered through a recombinant adeno-associated virus (rAAV) packaged as a self-complementary DNA. The notorious delivery limitations of rAAVs were overcome using a cross-correction approach that relied on the expression of the mutation in the context of glycosylphosphatidylinositoI-anchorless (ΔGPI) PrP. In this proof-of-concept study, we inoculated Rocky Mountain Laboratory (RML) prions into knock-in mice, in which the endogenous murine prion protein gene (*Prnp*) was replaced with the bank vole prion protein gene (Bv*Prnp*). Prion-inoculated mice that were retro-orbitally transduced with a protective rAAV vector encoding Bv*Prnp^V127^*ΔGPI survived ∼50 days longer than control mice that were unprotected. A deep proteomic analysis revealed that Bv*Prnp*^V127^ΔGPI was protective by slowing perturbations to the proteome observed in late-stage RML prion disease. In addition to capturing details of synaptic decay and depletion of proteins in proximity to PrP, the proteomic dataset revealed the identity of proteins of potential diagnostic value that may be central to the brain’s attempt to fight prion disease by contributing to astrocytosis or microgliosis, by coping with calcium influx, or by enhancing the endoplasmic reticulum processing of essential proteins. Taken together, our results demonstrate that a gene therapy based on a GPI-anchorless PrP containing the G127V mutation can delay the onset of prion disease in mice, providing a framework for development of a corresponding therapy in humans.

**AUTHOR SUMMARY:** A rare change in the human prion protein, involving a single building block, has been linked to strong protection against prion diseases—fatal neurodegenerative disorders. This study tested whether that protective effect could be reproduced using gene therapy in mice. To this end, we exposed the animals to infectious prions and then delivered the protective version of the protein into mice using a viral carrier. Treated mice survived about seven weeks longer than untreated animals, showing that the approach can meaningfully slow disease progression. To understand why, we examined changes in brain proteins during disease and found that treatment helped preserve the normal protein levels of cellular proteins, particularly those involved in communication between nerve cells. The analysis also identified proteins altered in the disease that are linked to the brain’s defense responses, including inflammation, stress handling, and protein processing, some of which may serve as future disease markers. Importantly, the limited protection observed was not due to poor delivery of the therapy but likely reflects biological limits of the model used. Overall, the findings support the idea that gene therapies based on naturally protective human variants could help slow prion diseases and improve understanding of how the brain responds to them.

## INTRODUCTION

Prion diseases are neurodegenerative diseases that afflict humans and a subset of mammals ^1^. To date, human manifestations of these diseases have remained incurable and are the cause of approximately 1 in 5,000 deaths ^2^. In these diseases, the cellular prion protein (PrP^C^), a small GPI-anchored protein ^3^ that is expressed from the prion protein gene (*Prnp*) in most vertebrate cells, converts through templated polymerization into a β-sheet-rich alternative conformer, referred to as PrP Scrapie (PrP^Sc^). In contrast to other neurodegenerative diseases whose etiologies are characterized by complex genetics contributing to disease risk, prion diseases are largely a one-gene disorder. Moreover, when working with prion-infected mice, these mice are not mere models for the disease but develop the disease with all its hallmarks, namely PrP^Sc^ deposition, astrogliosis, spongiform degeneration, and ultimately disease progression until death.

Several recent preclinical studies which evaluated PrP lowering approaches have generated optimism that an effective treatment can be found. Initially, a seminal proof-of-concept study of prion-infected mice, treated with antisense oligonucleotides (ASOs) designed to facilitate the destruction of *Prnp* transcripts and consequent lowering of PrP^C^ levels in the brain, documented dose-dependent survival extension in the absence of overt phenotypes ^4^. Despite this success, and evidence that the ASOs reached most brain cells, the study also revealed the limits of survival extension that may be achieved with this approach. When paired with the uncertainty of its translation into the clinic and the recurrent invasive intrathecal delivery required for ASOs, a need arises to continue exploring alternative therapy angles, including strategies based on rAAV vectors that deliver PrP^C^-lowering gene therapies. Amongst them are rAAV vectors coding for innovative payloads that silence prion gene expression by (i) directing compact epigenetic editors to the prion gene ^5^, (ii) instructing base editors to ablate prion gene transcription through the introduction of nonsense codons ^6^ and (iii) deploying zinc finger repressors (ZFRs) tailored to block prion gene transcription ^7^. All the above gene therapies have been reported to lower brain-wide PrP^C^ levels by more than 50%.

The ZFR-based strategy stands out because it was effective *in vivo* when delivered using PHP.B capsids as late as 120 days after the mice had been inoculated with prions, a time-point coinciding with the presentation of first overt prion disease symptoms in mice. Moreover, to overcome challenges with the delivery of rAAVs to human brain cells, the authors identified and made use of a capsid (STAC-BBB) whose brain tropism exceeded the corresponding tropism of AAV9 700-fold upon intravenous administration to cynomolgus monkeys. When administered to cynomolgus monkeys, the STAC-BBB-ZFR therapy achieved unprecedented reduction in PrP^C^ levels in a non-human primate after single intravenous dosing. A close look at the results from this study point toward two challenges. First, despite impressive performance characteristics of STAC-BBB, its PrP^C^ lowering capacity in cynomolgus monkeys was inferior to the corresponding potency of the PHP.B capsid used in mice. Second, although the administration of AAV-ZFRs lowered *Prnp* mRNA levels by >95% *in vitro* and to near undetectable levels within individual mouse neurons *in vivo*, all prion-inoculated mice, including those still alive at the 480-day study endpoint, exhibited weight loss and increased levels of the neurodegeneration biomarker Nfl, indicating that prion disease was not stopped altogether. The latter limitation indicates that therapies which merely lower PrP^C^ (as opposed to fully eliminating its expression) may not manifest as a cure due to residual PrP^C^ enabling prion replication and neurodegeneration. Consequently, there is a need to explore orthogonal gene therapy approaches that require the transduction of a lower percentage of brain cells for them to disarm the toxicity of PrP^Sc^.

The human PrP^V127^ mutation, discovered in the geographic epicenter of the kuru endemic in Papua New Guinea, may offer an untapped opportunity in this context. It has been credited with the survival of dozens of individuals among the Fore people who had been exposed to kuru disease through the practice of ritualistic cannibalism ^8^. Subsequent experimental work in mice, which were made co-transgenic for the human wild-type *PRNP* and mutant *PRNP^V12^*^7^ genes, established that the V127 mutation confers protection against several human prion inocula even when co-expressed at three-fold lower levels alongside wild-type human PrP^C^ ^9^. Thus, if a rAAV vector would instruct brain cells to produce the V127 mutant, the level of transduction required to achieve complete protection from prion disease may be lower than what would be required for equivalent protection through a treatment approach that aims to silence PrP expression.

In prior transgenic work that validated the protective capacity of PrP^V127^, both wild-type PrP and PrP^V127^ were co-expressed at consistent ratios in all cells responsive to the *Prnp* promoter because their expression was driven by the germ-line integration of the transgenes ^9^. In contrast, after virus transduction only a subset of cells will be positive for the rAAV vector-delivered transgene expressing PrP^V127^ and others will express none. Considering ways to address this potential shortcoming led us to cross-correction, a concept and mechanism that has had traction in the context of lysosomal storage disorders ^10^. In cross-correction, a small number of cells are turned into factories that release a therapeutic agent. Thereby, the therapeutic agent can act not only on the cells that produce it but also on neighboring cells. We hypothesized that cross-correction can be implemented for prion diseases by expressing a protective PrP^V127^ mutant lacking the GPI-anchor that normally attaches PrP^C^ to the outer leaflet of the cell membrane. This can be achieved by removing the sequence encoding the C-terminal GPI-anchor signal sequence from the prion gene ^11^.

It has been observed repeatedly that the lack of a GPI-anchor causes nascent PrP^C^ to become less N-glycosylated during its passage through the secretory pathway ^11^. Evidence suggests that both the PrP GPI-anchor and the N-glycans can pose steric hindrances to spontaneous and templated conversion into PrP^Sc^. Indeed, in transgenic mice over-producing a GPI-anchorless, underglycosylated PrP variant, PrPΔGPI spontaneously aggregates and deposits within the brain, causing clinical disease that is accelerated when membrane-anchored PrP^C^ is also present ^12, 13^. Anchorless PrP^C^ has also been shown to undergo templated *in vitro* conversion reactions more readily ^14^ and was observed to form denser deposits than wild-type PrP^C^ ^11^. Similarly, the recruitment of PrP^C^ into the templated polymerization can be hindered by N-glycans—a phenomenon that has been largely attributed to the negative charges contributed by the natural sialylation of N-glycans attached to PrP^C^ ^15^. In converse, PrP^C^ lacking N-glycans avoids these constraints and serves as a malleable substrate that exhibits a low seeding barrier in the templated conversion of a wide range of prion strains ^15^. Yet, as shown for RML strains, glycan-deficient recombinant and glycan-comprising brain-derived prion assemblies can give rise to identical parallel in-register intermolecular β-sheet (PIRIBS) structures ^16^. Finally, it has been reported that the half-life of anchorless PrP^C^ is six-fold longer than the corresponding value for GPI-anchored PrP^C^ ^17^; if this observation was to translate to brain-expressed protein, then it could boost the relative steady-state levels of anchorless PrP^C^. Although these characteristics of anchorless PrP^C^ would be undesirable when applied to the wild-type PrP^C^ sequence, as they would increase the risk of it acting as a substrate for spontaneous or templated polymerization, we hypothesized that in the context of anchorless PrP^V127^ they may turn into an advantage by helping the protective qualities of PrP^V127^ against prion diseases to manifest and by rendering it more effective against a wide range of prion strains.

Here, we began to put these ideas and concepts to the test in mice. In addition to asking if rAAV vectors that deliver the protective PrP^V127^ΔGPI mutant provide any protection, we characterized effects of RML prion disease on the brain proteome at an unprecedented level of depth. Although many of our observations were consistent with our expectations, others were not.

## RESULTS

### Design of an all-in-one rAAV vector of low immunogenicity that can promote cross-correction

Although the precise mechanism by which V127 confers protection is still being debated, there is broad agreement that the protective mutant acquires the same fold as native PrP^C^, barring minor structural differences surrounding the mutated residue ^18–21^. Similarly, when PrP^C^ is expressed without its N-glycans, a natural consequence of the removal of the GPI-anchor attachment sequence, it will still acquire its natural PrP-fold ^22–24^ (**Fig 1A**). These insights were important for immunological considerations, as they predicted that the expressed therapeutic agent would have low immunogenicity. In contrast, rAAV capsids can be expected to illicit a moderate immune response in naïve mice. Whereas in humans prior exposure to natural AAVs represents a major consideration ^25^, for studies in laboratory mice this concern is small if no repeat administrations are planned. Consequently, so long as the design of the synthetic coding sequence for the transgenic expression of PrP^V127^ avoided highly immunogenic CpG islets ^26^, the packaged therapeutic construct was expected to exhibit low immunogenicity, largely restricted to the DNA payload. To mitigate immunogenicity of the latter, we capitalized on redundancies in the genetic code and eliminated most occurrences of CpGs (**Fig 1B**).

**Fig 1.**
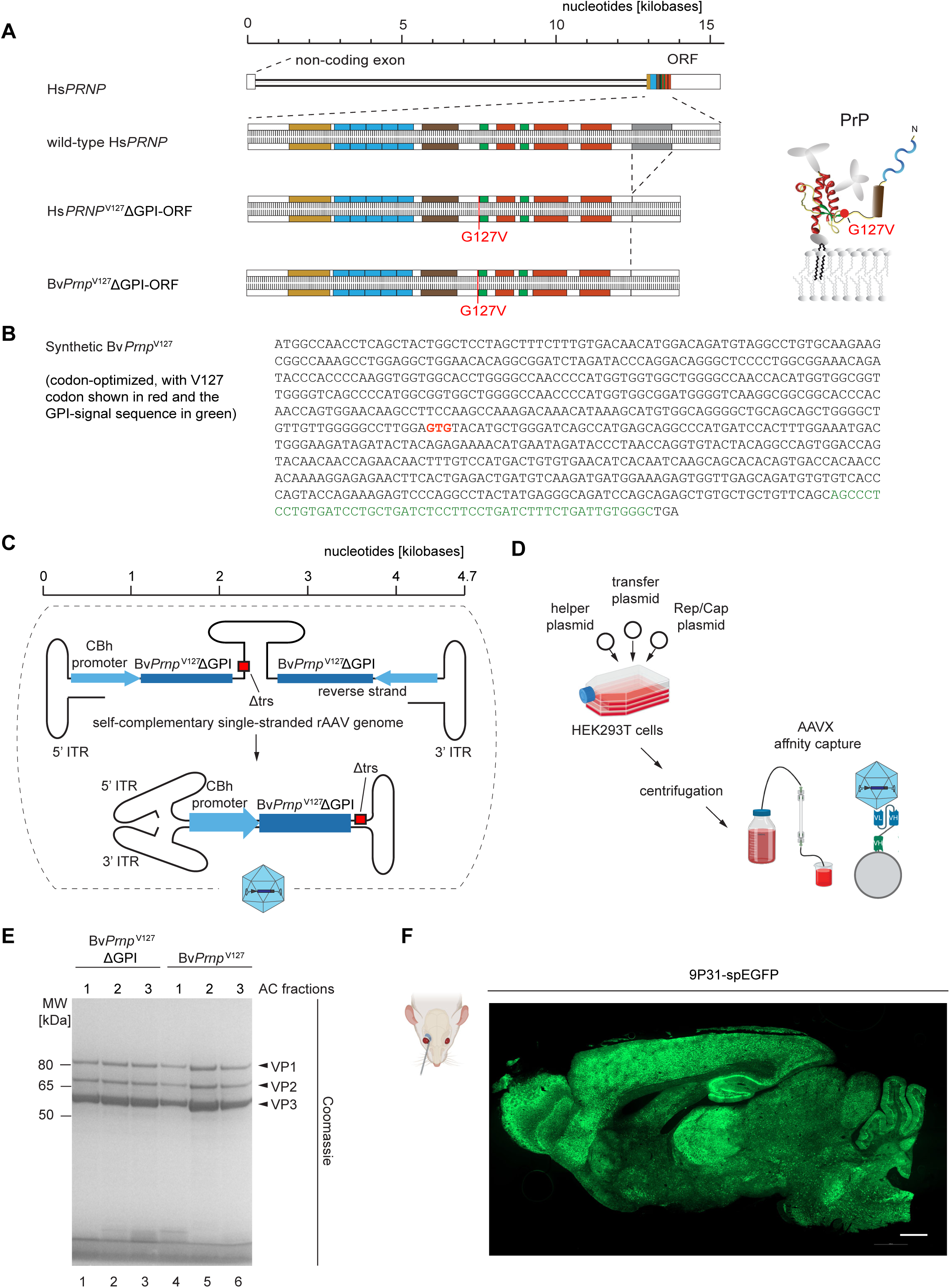
Design of an all-in-one rAAV vector of low immunogenicity that can promote cross-correction. (A) Genome organization of the human (Hs) *PRNP* gene. Zoom-in depicts the sequence organization of the ORF. Colors highlight coding segments as follows: beige, signal peptide for co-translational ER targeting; blue, octarepeats; brown, hydrophobic region; green, β-sheets; red, α-helices; grey, signal sequence for attachment of GPI-anchor. The protein structure model shown on the right identifies the approximate position of the G127V residue within the tertiary fold of PrP^C^ with a red circle. For ease of orientation, the image depicts a PrP^C^ molecule that is inserted into a membrane bilayer with a GPI-anchor. (B) Nucleotide sequence of a codon-optimized, synthetic bank vole *Prnp*^V127^ coding sequence. The V127 codon and the GPI-SS are shown in red and green font, respectively. (C) Design of self-complementary rAAV vector for the expression of synthetic bank vole PrP^V127^ΔGPI. (D) Schematic summarizing key steps of rAAV purification method based on assembly of virus in Hyperflasks and AAVX affinity capture purification. (E) Assessment of purity and yield of representative rAAV vector preparations coding for wild-type Bv*Prnp*^V127^ΔGPI or Bv*Prnp*^V127^ using SDS-PAGE followed by Coomassie blue staining. VP1, VP2 and VP3 designate the three viral proteins that constitute the viral capsid in an expected 1:1:10 relative abundance ratio. (F) Representative image showing pronounced expression and brain-wide distribution of spEGFP signal in sagittal cut of mouse brain following retro-orbital administration of 9P31-spEGFP vector. Scale bar = 100 µm.

Rather than working with wild-type mice, we selected for this study the recently introduced bank vole knock-in (Bv*Prnp* ki) model in which the endogenous mouse *Prnp* open reading frame (ORF) was replaced with the respective Bv*Prnp* ORF encoding isoleucine at polymorphic codon 109 ^27^. Two considerations guided this choice: 1) Bv*Prnp*-expressing mice had been shown to develop disease faster than wild-type mice ^28^. 2) In anticipation of future inoculation work with human prions, we hoped to capitalize on bank vole PrP’s universal prion acceptor characteristics, when we translate results from work with mouse prion inocula (this work) to future work with human prion inocula ^29, 30^.

To maximize the expression of the heterologous Bv*Prnp*^V127^ΔGPI, the payload was designed as a self-complementary DNA, thereby avoiding the replication step that single-stranded AAV vectors must undertake upon transduction ^31^, a limiting biology dependent on the availability of host cell factors (**Fig 1C**). Specifically, the expression cassette of the construct was flanked by inverted terminal repeats (ITRs) derived from the AAV2 serotype, with one of the ITRs carrying a small deletion within the terminal repeat sequence (TRS) motif known to manifest in self-complementarity ^32^. The expression of Bv*Prnp*^V127^ΔGPI was driven by the chimeric chicken β-actin promoter with intron (CBh) that had been shown to promote the steady expression of transgenes in mouse brains for extended periods ^33^.

While the study was ongoing, several innovations emerged in the rAAV vector field, including alternative ways to purify rAAV vectors as well as capsids with optimized blood brain barrier (BBB) penetrance. Consistent with the pilot format of the study, we incorporated comparisons of two of these tool developments in the study design: 1) an affinity capture-based method for rAAV vector purification using AAVX matrices, which we compared to the more widely used iodixanol gradient centrifugation (**Fig 1D**) ^34^, and 2) a capsid, designated 9P31 (Voyager Therapeutics Inc.), which was reported to feature several-fold higher CNS penetrance than PHP.eB, which is often the choice capsid for targeting the CNS in mice following intravenous administration ^35^. Purity assessments of rAAV vectors that we prepared by AAVX affinity chromatography showed predominantly three bands derived from the three capsid isoforms VP1, VP2, and VP3 in the expected approximate ratio of 1:1:10 after SDS-PAGE separation (**Fig 1E**). To test the 9P31 vector, we assembled a self-complementary rAAV vector identical to the one described for the expression of Bv*Prnp*^V127^ΔGPI, except that the Bv*Prnp*^V127^ΔGPI ORF was replaced with a coding sequence for a variant of the enhanced green fluorescent protein, which we directed to the endoplasmic reticulum (ER) by the addition of a signal peptide (spEGFP, with the signal peptide-encoding sequence borrowed from the calreticulin gene) (**Fig 1F**) ^36^. As results from our subsequent experiments will show, the combination of the 9P31 capsid with the AAVX purification method gave rise to the highest expression levels of heterologous constructs.

### Retro-orbital injection of BvPrnpG127VΔGPI into prion-inoculated mice caused similar survival extension regardless of rAAV vector preparation method or capsid choice

As we were gearing up for the study, our breeding stock of Bv*Prnp* ki mice produced relatively small litters, which forced us to replace plans of parallel inoculations with staggered inoculations. We hypothesized that the expression of V127 within the context of an anchorless Bv*Prnp* expression construct (Bv*Prnp*^V127^ΔGPI) might lead to the longest survival extension due to enhanced cross correction. Consequently, we began the *in vivo* studies by comparing side-by-side mice that had been prion-inoculated on the same day, then were either left untransduced or were retro-orbitally transduced 60 days later with rAAV vectors coding for s*pEGFP* (negative controls) or Bv*Prnp*^V127^ΔGPI (**Fig 2A**). The latter cohort consisted of three sub-cohorts. Two of these were transduced with 9P31 vectors that had been purified either by iodixanol density centrifugation or AAVX affinity chromatography (**Fig 1D**). The third sub-cohort was transduced with rAAV vectors encapsulated in the PHP.eB capsid that had also been purified by AAVX affinity chromatography. Deferred were the analyses of mice transduced with 9P31 encapsulated vectors coding for anchored BvPrP^V127^.

**Fig 2.**
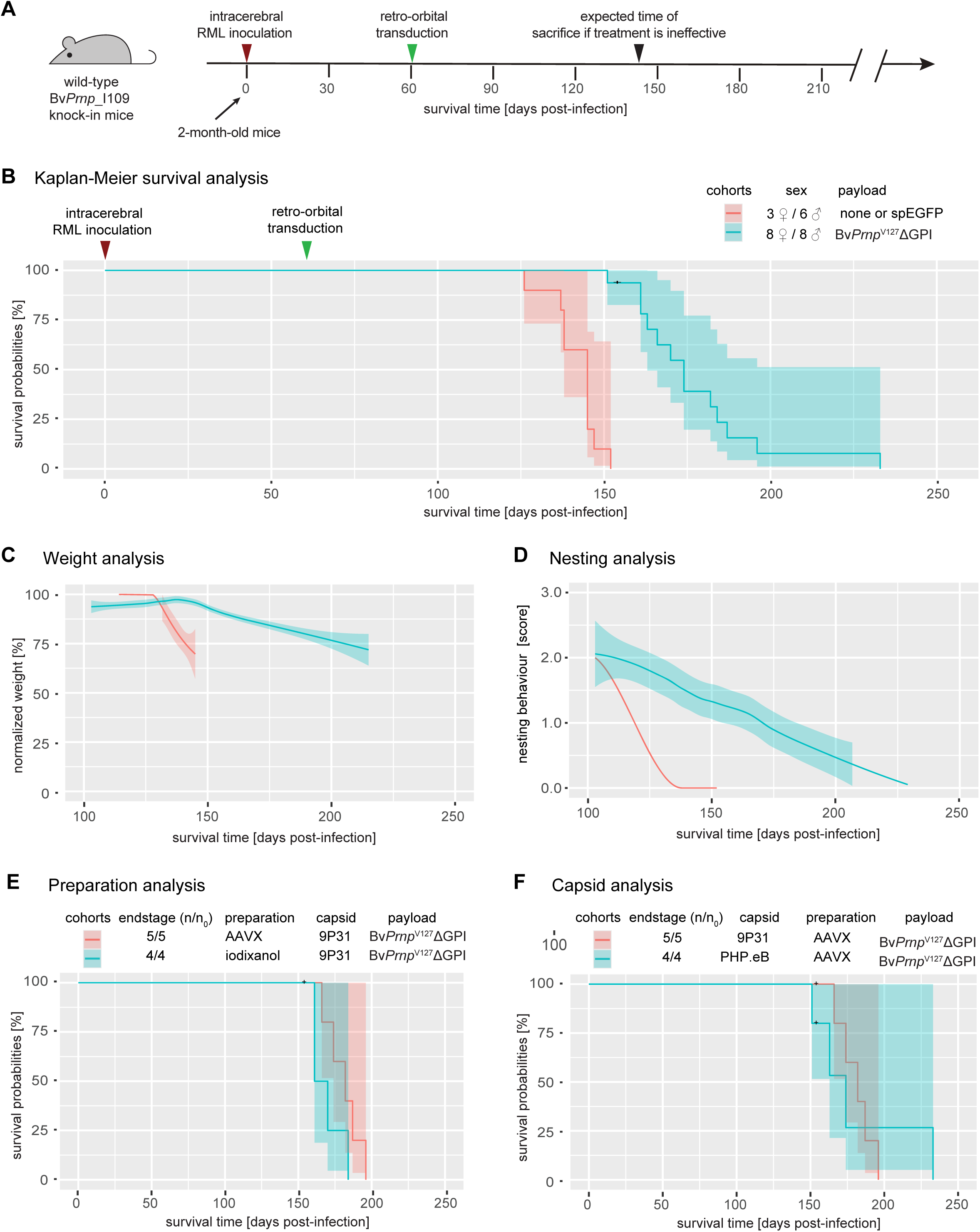
Heterologous expression of rAAV vector-delivered BvPrP^V127^ΔGPI causes survival extension in RML-infected *BvPrnp ki* mice. (A) Timeline of pilot *in vivo* prion infection and treatment study in *BvPrnp* ki mice. Note that all endogenous and heterologous bank vole sequences employed in this work carried the I109 polymorphism. (B) Kaplan-Meier chart comparing the survival of negative control mice and treated mice. (C) Weight analysis chart depicts the relative weights of mice shown in Panel B from the time of 100 dpi to the time of sacrifice when the animals met humane prion disease endpoints. Note that weights of individual mice were normalized to their weight at 100 dpi to account for individual weight differences and sex-specific weight differences. (D) Nesting analysis chart showing the nesting score of mice shown in Panel B from 100 dpi to the end of the study. (E) Chart documenting that the alternative rAAV vector preparation methods based on AAVX affinity capture or iodixanol gradient centrifugation for purifying the therapeutic 9P31 - *BvPrnp*^V127^ΔGPI had an insignificant influence on survival extension. (F) No significant differences in the survival times afforded by the expression of *BvPrnp*^V127^ΔGPI was observed whether the therapeutic payload was encapsulated in the PHP.eB or 9P31 capsid. In all graphs, 95% confidence intervals are indicated by shading around the curves.

As expected, RML-prion inoculated Bv*Prnp* ki mice began to show symptoms 130-150 days post-inoculation (dpi) and had to be sacrificed shortly thereafter when they reached prion disease endpoint (**Fig 2B**). Whether the prion-inoculated mice were left untransduced or transduced with the negative control 9P31-*spEGFP* vector made no significant difference to their survival. In contrast, the cohort of mice that had been made to express the therapeutic Bv*Prnp*^V127^ΔGPI construct, survived approximately 50 days longer. One characteristic of the Kaplan-Meier curve for this cohort is a relatively broad spread of survival times from 150 to a maximum of 232 days, possibly reflecting variances associated with the administration of the therapeutic capsids (**Fig 2B**). Plotted results from monitoring body weights (**Fig 2C**) and nesting scores (**Fig 2D**) of the mice reflected both the survival extension as well as the more gradual decline in the Bv*Prnp*^V127^-ΔGPI cohort. When the latter results were deconvoluted based on how the 9P31-Bv*Prnp*^V127^ΔGPI capsids were prepared (**Fig 2E**) or whether the payload was encapsulated in 9P31 or PHP.eB, subtle shifts to longer survival time were observed when rAAV vectors had been AAVX purified and encapsulated in 9P31, but no significant differences emerged (**Fig 2F**).

### Survival extension afforded by Bv*Prnp*^V127^ΔGPI is accompanied by reduced PrP^Sc^ accumulation but does not correlate linearly with steady-state expression levels of therapeutic payload

To begin to understand how the expression of Bv*Prnp*^V127^ΔGPI had delayed the disease, at 152 dpi, i.e., when the prion-inoculated control mice reached prion disease endpoint, a small number of Bv*Prnp* ki mice, which had been treated with the 9P31 encapsulated vector coding for the Bv*Prnp*^V127^ΔGPI mutant were also sacrificed, along with age-matched control Bv*Prnp* ki mice that had neither been prion-inoculated nor transduced. Sagittal half brains of these mice were formalin-fixed, and the remaining half brains were homogenized and extracted, then processed for western blot analyses with or without prior digestion with proteinase K (PK). This analysis revealed the expected increase in total PrP levels, a result of the accumulation of PrP^Sc^, in RML prion-inoculated mice (relative to uninoculated age-matched control mice) that had been transduced with 9P31-*spEGFP* negative control viruses (**Fig 3A**). In contrast, RML prion-inoculated mice that had been transduced with the therapeutic 9P31-Bv*Prnp^V127^*ΔGPI vector showed lower levels of PrP^Sc^ accumulation but were not devoid of PK-resistant PrP^Sc^ (**Fig 3B**).

**Fig 3.**
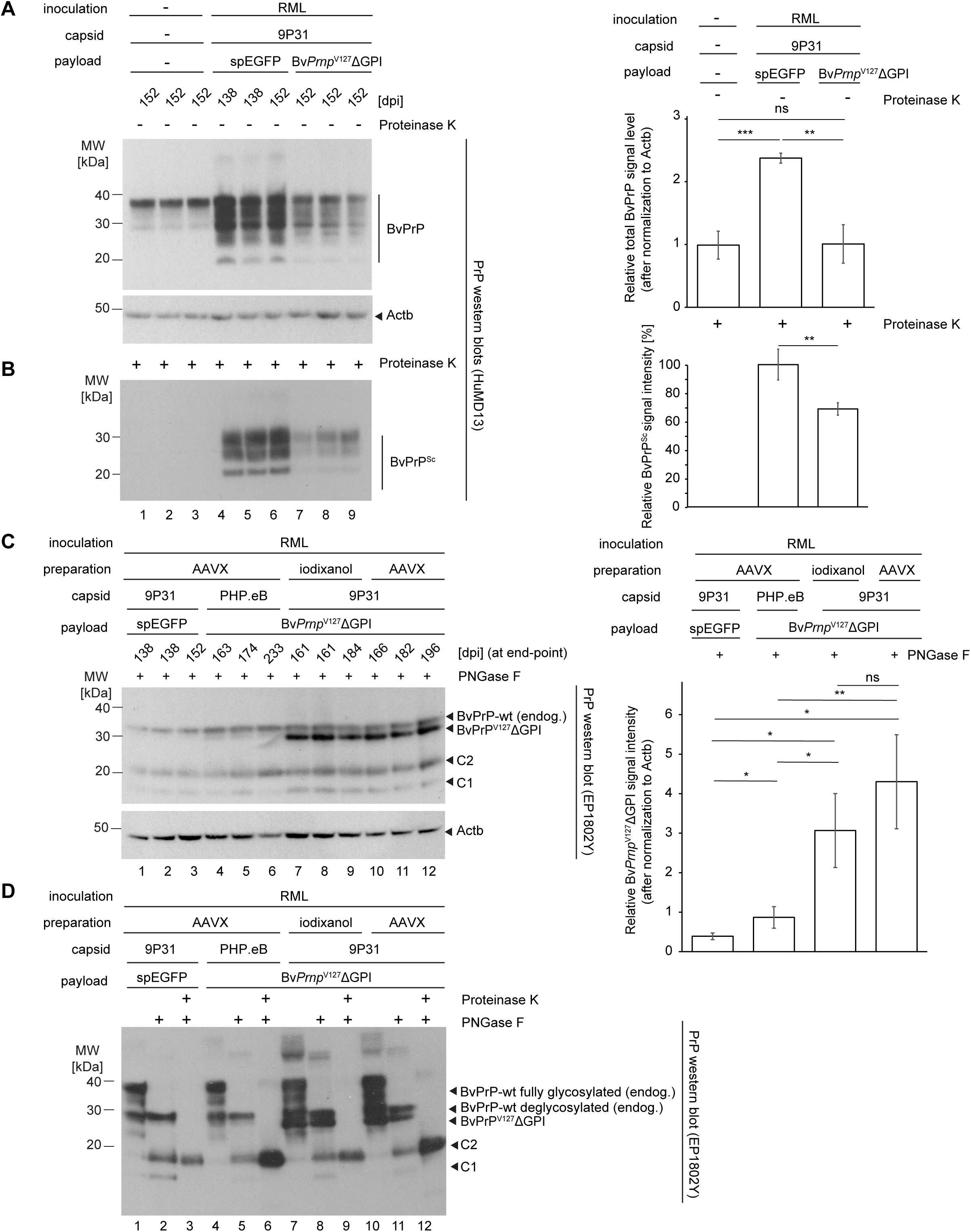
Expression of heterologous Bv*Prnp*^V127^ΔGPI in RML-inoculated *BvPrnp* ki mice mitigates accumulation of PrP^Sc^ and increases total PrP signal intensities up to threefold. (A) Western blot analysis of extracts from sagittal half brains of Bv*Prnp* ki mice that were either from a cohort of naïve mice, or RML-inoculated mice that were mock-treated with a 9P31-spEGFP vector or treated with the 9P31-Bv*Prnp*^V127^ΔGPI vector. Note that all animals were sacrificed by 152 days, i.e., when the RML-inoculated, mock-treated cohort reached its prion disease endpoint. Neither the naïve *BvPrnp* ki cohort nor the mice treated with 9P31-Bv*Prnp*^V127^ΔGPI showed overt signs of prion disease at the time of their sacrifice. A western blot depicting steady-state beta actin (Actb) levels served as a loading control in this analysis. The bar graph on the right revealed significant differences (Student’s t-test) in total PrP levels in the three cohorts. (B) Analysis of proteinase K-resistant PrP^Sc^ in protein extracts analyzed in Panel A. The bar graph on the right quantified band intensities of proteinase K-resistant PrP^Sc^. It was normalized to PrP^Sc^ levels in the mock-treated cohort transduced with the 9P31-spEGFP vector. (C) Therapeutic vectors encapsulated in 9P31 caused robust expression of Bv*Prnp*^V127^ΔGPI that exceeded endogenous total PrP levels threefold. Western blot analysis of extracts from sagittal half brains of Bv*Prnp* ki mice that were RML-inoculated but then were either mock-treated through transduction with the spEGFP vector or transduced with therapeutic vectors coding for Bv*Prnp*^V127^ΔGPI. For this analysis, brains from all mice were harvested after the animals reached their individual prion disease endpoint as shown by their unique dpi. To facilitate the recognition of the therapeutic Bv*Prnp*^V127^ΔGPI signal amongst a complex mixture of N-glycosylated endogenous BvPrP expression products, N-glycans were removed by PNGase F digestion prior to the western blot analysis. Note the pronounced signal of the Bv*Prnp*^V127^ΔGPI band, which migrated as expected a bit faster than the endogenous wild-type full-length BvPrP band on account of it lacking the GPI-anchor. The detection of Actb served again as loading control in this analysis. The bar graph on the right quantifies the intensities of Bv*Prnp*^V127^ΔGPI signals in the treatment cohorts. (D) Side-by-side analysis of one brain extract from each of the four treatment cohorts analyzed in Panel C with or without digestion with Proteinase K followed by digestion with PNGase F. Note that the bands visible in lanes 6, 9, and 12 don’t appear to constitute doublet signals, which would be expected if the anchorless BvPrP^V127^ΔGPI had contributed to the Proteinase K-resistant PrP^Sc^.

Once the remaining cohort of the 9P31-Bv*Prnp^V127^*ΔGPI-treated mice had reached the disease endpoint, we undertook additional western blot analyses, after processing their half brains as above. These experiments were guided by an interest in understanding why the sub-cohort, which had been treated with the PHP.eB-encapsulated payload coding for Bv*Prnp*^V127^ΔGPI had shown a similar survival extension as the mice whose identical therapeutic construct had been encapsulated in the 9P31 capsid, a counterintuitive result based on our prior observation that 9P31 vectors mediated an approximately seven-fold higher CNS transduction than PHP.eB vectors when payloads, virus preparation, and administration steps were identical ^37^. Since we were particularly interested in determining the steady-state Bv*Prnp*^V127^ΔGPI expression levels that had been reached in the treated cohort, for this western blot analysis the total brain extracts were first digested with PNGase F so that PrP isoforms differing solely in their N-linked glycans would be reduced to a single band (**Fig 3C**). Intriguingly, this analysis corroborated the relative CNS transduction potencies of 9P31 and PHP.eB by showing that the Bv*Prnp*^V127^ΔGPI-derived western blot signals were considerably stronger when the payload had been encapsulated in 9P31 than the respective signal in brain extracts of mice that had been transduced with the corresponding PHP.eB vector. In fact, the western blot signal that we interpreted to represent Bv*Prnp*^V127^ΔGPI by its absence in non-transduced RML-inoculated mice or in mice that had been transduced with 9P31-*spEGFP* emerged as the strongest PrP antibody-reactive signal. The intensity of this band in 9P31-Bv*Prnp^V127^*ΔGPI mice exceeded approximately threefold the respective signals for endogenous wild-type BvPrP. This result indicated that the transduction worked better than we had anticipated but it also indicated that the therapeutic potency of Bv*Prnp*^V127^ΔGPI, when measured based on the survival extension it conferred, did not correlate linearly to its expression level in Bv*Prnp* ki mice.

Finally, we sought to determine if the highly expressed BvPrP^V127^ΔGPI contributed to the formation of Proteinase K resistant material in the treatment cohort after these mice had succumbed to prion disease. To reduce the complexity of signals expected if isoforms with varying N-glycan occupancy were present, we first digested representative brain extract from each of the four cohorts with Proteinase K, then removed N-glycans from the resolubilized PrP^Sc^ material by an additional digestion with PNGase F. The side-by-side analysis of the resultant PrP products next to undigested or only PNGase F digested samples revealed that mice, which expressed both the endogenous Bv*Prnp* gene and the synthetic Bv*Prnp*^V127^ΔGPI construct, still only gave rise to a single PrP^Sc^ signal which ran at the same level as the respective band in mice that were not transduced (**Fig 3D**). Previous work with prion-inoculated transgenic mice that expressed both wild-type PrP and anchorless PrPΔGPI documented that PK-resistant unglycosylated bands derived from anchorless PrPΔGPI run considerably faster than the corresponding wild-type-derived bands and can easily be distinguished in a western blot analysis ^11^. These results suggest that BvPrP^V127^ΔGPI did not contribute in RML-infected mice to the formation of PK-resistant PrP^Sc^, even though the mice expressed relatively high levels of the protective variant and died at ∼200 dpi of symptoms consistent with a late-stage prion disease diagnosis.

### Expression of Bv*Prnp^V127^*ΔGPI slows perturbations to the proteome observed in prion-inoculated mice

Although a fair bit is known about how prion diseases affect specific proteins, and it has been reported that the disease affects global protein synthesis following activation of the unfolded protein response ^38^, systematic in-depth proteome comparisons of age-matched naïve and prion-inoculated end-state prion disease brains have, to our knowledge, not been reported. Moreover, despite reports of V127 conferring some resilience to structural dynamics *in vitro* ^21, 39^, our understanding of how V127 confers its protection against prion diseases remains limited. Although it is apparent that the expression of V127 slows the accumulation of PrP^Sc^, it is not known if its protective effect manifests in a mere slowing of prion-disease-associated perturbations or involves more specific protective changes to the proteome. To fill these knowledge gaps, we sought to interrogate the proteome of brain extracts from age-matched naïve Bv*Prnp* ki mice, and the corresponding prion-inoculated and mock 9P31-spEGFP- versus 9P31-Bv*Prnp^V127^*ΔGPI-treated mice, i.e., the same samples we had subjected to western blot analysis (**Fig 3A**).

To prepare the samples for mass spectrometry analyses, the total brain extracts were fully denatured in the presence of urea, then reduced and alkylated, and finally trypsinized. Using 30-minute gradients, tryptic mixtures were separated on a reversed phase column. The effluents from this separation were online coupled by nanospray ionization to an Orbitrap Astral mass spectrometer, which was operated in data-independent acquisition (DIA) mode (**Fig 4A**). A high degree of run-to-run proteomic sequence coverage afforded by the DIA acquisition mode obviated the need for isobaric labeling, thereby enabling MS2-based relative quantitation of consecutively analyzed samples. Taken together, this configuration allowed the deep unsupervised characterization of the mouse brain proteome, leading to the relative quantification of 4,874 proteins in all cohorts, comprised of three biological replicates for each of the three cohorts (**Fig 4B**). More than 800 of these proteins were identified based on peptide-to-spectrum matches (PSMs) that accounted for more than 50% coverage of their protein sequence, with approximately half of the protein identifications supported by ≥100 PSMs, and more than 4,800 protein identifications based on ≥10 PSMs. Even at cursory inspection, it was apparent that the three biological replicates for each cohort were more like one another than samples from other cohorts, an impression supported by results from an unsupervised hierarchical clustering analysis (see below).

**Fig 4.**
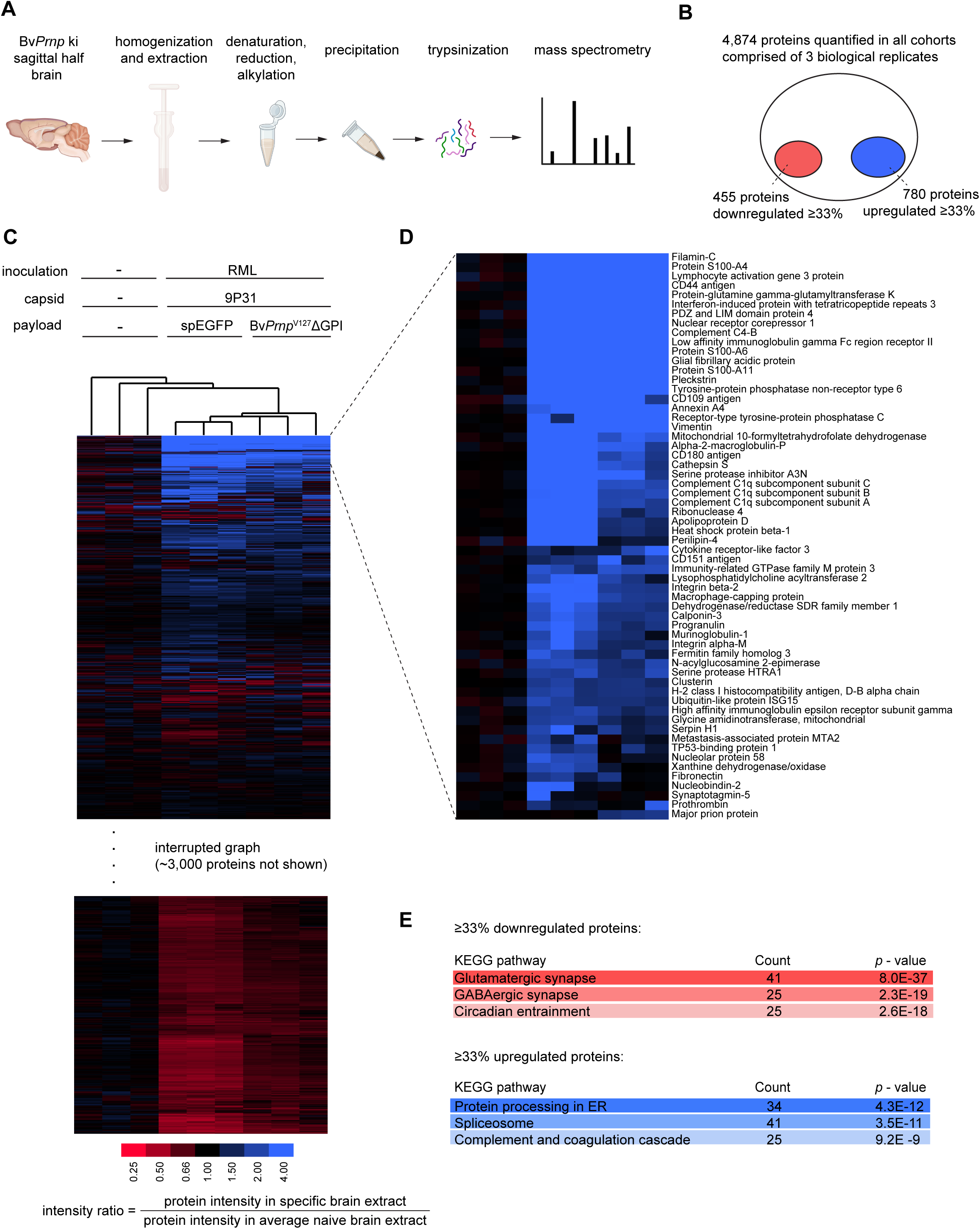
Expression of Bv*Prnp^V127^*ΔGPI slows perturbations to the proteome observed in prion-inoculated mice. (A) Schematic showing sample workup scheme for global proteome analysis. (B) Global proteome summary diagram documenting benchmarks of the analysis, including the number of proteins whose steady-state levels in RML-inoculated and mock-treated brains were ≥ 33% altered relative to their average protein levels in naïve mouse brains. (C) Result from hierarchical clustering analysis of global proteome dataset. Note that the unbiased clustering analysis grouped the cohorts in accordance with the study design, i.e., it identified three groups of highly similar brain extracts. The global proteomic signature of the mice, which were RML-inoculated and transduced with 9P31-delivered *BvPrnp*^V127^ΔGPI vectors, places them between the naïve and the RML-inoculated but mock-treated mouse cohort. (D) Zoom-in into the hierarchical clustering results focused on proteins whose steady-state levels are upregulated in RML-inoculated mice. The graph indicates highly consistent changes in the direction of perturbation for a given protein and corroborates that many proteins are to a lesser degree altered in their steady-state levels in the 9P31-Bv*Prnp*^V127^ΔGPI treatment cohort than the 9P31-spEGFP mock-treated cohort. (E) Short-listed results from KEGG pathway analyses undertaken with proteins whose levels were ≥ 33% up- or down-regulated.

When the relative abundances of individual proteins in all nine samples were computed by forming ratios of their combined ion intensities in each sample and their respective average ion intensities in the naïve Bv*Prnp* ki cohort, it emerged that most proteins were not altered by more than 33% in their steady-state levels in the disease. Yet, taking the opposite perspective, 780 genes were ≥33% upregulated in their expression and 455 proteins were ≥33% down-regulated at end-stage prion disease, relative to age-matched naïve Bv*Prnp* ki mice. Bv*Prnp*^V127^ΔGPI-treated mice fell between these two extremes and, importantly, exhibited no proteomic drifts relative to naïve mice that were not also observed in the RML-inoculated mice sacrificed at the humane prion disease endpoint. However, the abundance changes were less pronounced for these treated mice than what was observed for the respective protein abundances in the 9P31-spEGFP-transduced mice (**Fig 4C,D**). Taken together, these observations were consistent with the interpretation that the expression of Bv*Prnp^V127^*ΔGPI does not prolong survival by having a specific effect on a subproteome which can compensate for prion disease-induced proteome perturbations.

We next undertook pathway analyses to investigate more systematically the changes to the proteome that manifested in RML-inoculated Bv*Prnp* ki mice. To this end, we interrogated the KEGG pathway repository with identifiers of proteins that were either ≥33% upregulated or downregulated in the disease (**Fig 4E**). Consistent with expectations, these analyses revealed a highly significant reduction in the steady-state levels of proteins associated with synaptic biology. Interestingly, within the various types of synapses, glutamatergic synapses were the most significantly impacted (*p* = 8.0 E-37), with the GABAergic (2.3 E-19), adrenergic (1.5 E-13), dopaminergic (5.9 E-13), and cholinergic synapses (1.2 E-12), showing lesser impacts but still returning highly significant *p*-values. Consistent with known perturbations of prion diseases to sleep-wake patterns, the KEGG pathway defining components of circadian entrainment was also one of the most downregulated (*p* = 2.6 E-18).

Brain cells in RML-inoculated Bv*Prnp* ki mice impacted by PrP^Sc^ accumulation do not succumb without a fight, as several pathways can be seen to be ≥33% upregulated relative to age-matched naïve *BvPrnp* ki mice. Chief among them are the KEGG pathway defining components that facilitate ‘Protein processing in the endoplasmic reticulum’ (*p* = 4.3 E-12). Along with ER proteins, spliceosome proteins were next in this category in the order of relative significance (*p* = 3.5 E-11). Taken together, these analyses added granularity to the notions that prion diseases are characterized by marked deficiencies in synaptic biology and synaptic entrainment. The latter manifest alongside attempts of brain cells to delay their demise by increasing their ER processing capacity.

### Threefold increase in total PrP and 20-fold higher levels of unglycosylated PrP in mice transduced with *BvPrnp^G127V^*-ΔGPI at gene levels equivalent to endogenous Bv*Prnp*

Next, we sought to assess the extent to which the brain-wide delivery of the genetic payload of our gene therapy had occurred. To this end, we capitalized on nucleotide sequence differences in the endogenous Bv*Prnp* ORF and the synthetic Bv*Prnp*^V127^ΔGPI-ORF. Specifically, we identified a stretch within the respective ORFs that could be amplified with the same PCR primers because it was flanked by identical nucleotide sequences yet could only be digested by one or another restriction enzyme, namely EagI for endogenous Bv*Prnp* gene sequences and CsiI for Bv*Prnp*^V127^ΔGPI inserted into the genome following transduction (**Fig 5A**). By comparing the signal intensity of the fragments released by the respective restriction enzymes, this assay could inform about the relative amount of genomic DNA (gDNA) coding for BvPrP that was contributed by the endogenous and heterologous synthetic ORF. This analysis revealed that the relative amounts of genomic endogenous Bv*Prnp* ORFs were approximately matched to those of the heterologous synthetic Bv*Prnp*^V127^ΔGPI ORFs in mice that had been transduced with the 9P31- Bv*Prnp*^V127^ΔGPI vectors (**Fig 5B**).

**Fig 5.**
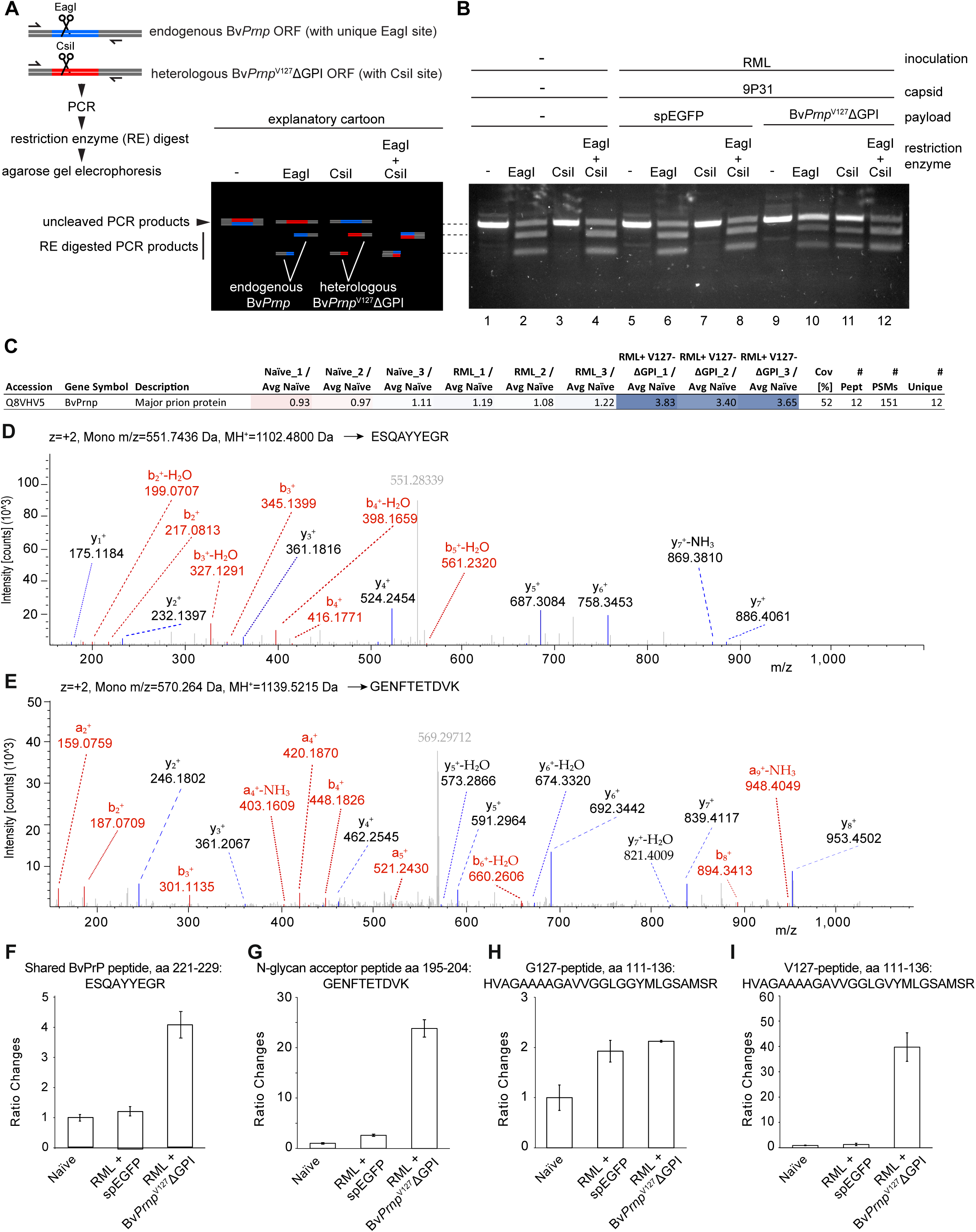
Threefold increase in total PrP and 20-fold higher levels of unglycosylated PrP in mice transduced with *BvPrnp^G127V^*-ΔGPI at gene levels equivalent to endogenous Bv*Prnp*. (A) Schematic of PCR and restriction enzyme (RE) digest analysis of genomic brain DNA preparations of mice for determining the relative abundance of endogenous and heterologous *BvPrnp* ORFs. (B) Transduction of mice with 9P31-Bv*Prnp*^V127^ΔGPI gave rise to the detection of RE cleavage products specific for heterologous Bv*Prnp*^V127^ΔGPI. Note that the signal intensities of RE digested PCR products in lanes 10 and 11 are approximately matched, indicating similar levels of *BvPrnp* and Bv*Prnp*^V127^ΔGPI ORFs in the genome. (C) Steady-state total BvPrP levels, comprising both endogenous and heterologous BvPrP expression products, revealed in global proteomic analysis based on 52% sequence coverage and the assignment of 151 PSMs to 12 unique BvPrP-derived peptides. (D) MS2 spectrum of the tryptic peptide ESQAYYEGR contributed by endogenous and heterologous pools of BvPrP. (E) MS2 spectrum of the tryptic peptide GENFTETDVK comprising the second ‘NxT’ N-glycan acceptor consensus motif within BvPrP. (F-I) Quantitation of tryptic BvPrP peptides that are either present in all BvPrP sequences analyzed (F), only observed in the sub pool of BvPrP that is non-glycosylated at GENFTETDVK (G), only present in endogenous Bv*Prnp* expression products (H), or only present in the heterologous Bv*Prnp*^V127^ΔGPI expression product (I).

The depth of the proteome dataset that the data-independent acquisition mode afforded allowed additional questions to be answered by comparing the total levels of BvPrP observed in the respective brain extracts to the levels at which specific BvPrP-derived peptides were detected. Consistent with the western blot data and the genomic DNA analyses, the mass spectrometry-based quantitation indicated that the total steady-state BvPrP in mice which had been transduced with the 9P31-Bv*Prnp*^V127^ΔGPI-ORF vector was approximately three-fold higher than the respective quantity in the naïve Bv*Prnp* ki mice (**Fig 5C**). We next investigated the levels of three specific BvPrP-derived tryptic peptides in the nine samples, namely 1) the peptide ESQAYYEGR present in both endogenous and heterologous PrP (**Fig 5D**), 2) the peptide GENFTETDVK, harboring one of the two N-linked glycan acceptor sites (NxT) present in the protein (**Fig 5E**), and 3) the respective BvPrP peptides that surround the G127 or V127 residues and therefore can be unequivocally assigned to endogenous BvPrP or the heterologous BvPrP^V127^ΔGPI, respectively. These analyses detected the generic BvPrP peptide at lowest levels in the naïve mice, at slightly increased levels in RML-inoculated mice, and at approximately three-fold higher levels in the 9P31-Bv*Prnp*^V127^ΔGPI-ORF-transduced mice (**Fig 5F**). The peptide harboring the unmodified N-glycan acceptor site was present at approximately 20-fold higher levels in the Bv*Prnp*^V127^ΔGPI-ORF-transduced mice. Considering the three-fold higher total BvPrP levels in these mice, this result suggests that the heterologous anchorless protein is 6- to 7-fold less likely to carry the N-glycan at the respective acceptor site than the endogenous protein (**Fig 5G**). The BvPrP peptide comprising the wild-type BvPrP G127 residue was observed at about twice the level in the RML-inoculated samples (**Fig 5H**), consistent with the stabilization and overall increase in steady-state PrP levels that are commonly observed in prion-infected mice. Finally, as anticipated, the V127-comprising peptide was not detected in naïve or RML inoculated mice transduced with the control 9P31-spEGFP vector but was robustly detected in mice transduced with 9P31-Bv*Prnp*^V127^ΔGPI (**Fig 5I**).

These results corroborated the increase in total BvPrP levels, which we had observed by western blot analyses (**Fig 3**). However, these data also indicated that this increase at the protein level was not reflected at the gDNA level, where endogenous and heterologous Bv*Prnp* ORFs were approximately matched in 9P31-Bv*Prnp*^V127^ΔGPI transduced brains. This divergence of steady-state protein levels versus gene copy number may reflect differences in the relative strengths of the endogenous mouse *Prnp* promoter versus the CBh promoter used to drive the heterologous expression. Alternatively, these data may indicate differences in the half-lives of anchored versus secreted 9P31-Bv*Prnp*^V127^ΔGPI. The results further validated the anticipation that the expression of anchorless Bv*Prnp*^V127^ΔGPI is accompanied by a marked reduction in the N-glycan occupancy of the heterologous protein relative to endogenous anchored BvPrP.

### Bv*Pr*npV127ΔGPI reduced levels of PK-resistant PrP^Sc^ relative to anchored *BvPrnp^V127^*

To begin to assess if cross-correction of an anchorless Bv*Prnp*^V127^ΔGPI expression construct can supersede the protection conveyed by the expression of anchored Bv*Prnp*^V127^, we next compared directly the survival curves of Bv*Prnp* ki mice that had been transduced with 9P31-delivered virus vectors, whose therapeutic payload differed only in the inclusion or omission of the GPI-attachment sequence within the protective Bv*Prnp*^V127^ ORF. Consistent with the hypothesis that cross-correction may enhance the protective capacity of Bv*Prnp*^V127^ΔGPI transduced mice, the Kaplan-Meier curves of mice that had received retro-orbital injections of the 9P31-delivered vectors coding for anchored Bv*Prnp*^V127^ led to a shorter survival extension of ∼25 days (**Fig 6A**). This finding was validated by western blot analyses of brain extracts of age-matched mice from the two treatment cohorts. Specifically, these analyses documented that levels of PK-resistant PrP^Sc^ were significantly higher (30%, p<0.05) in mice that had been treated with the expression construct coding for anchored V127 (**Fig 6B**). As in our previous western blot analyses (**Fig 3**), no evidence of PK-resistant Bv*Prnp*^V127^ΔGPI was detected in these western blots when we analyzed samples of 9P31-Bv*Prnp*^V127^ΔGPI-transduced mice.

**Fig 6.**
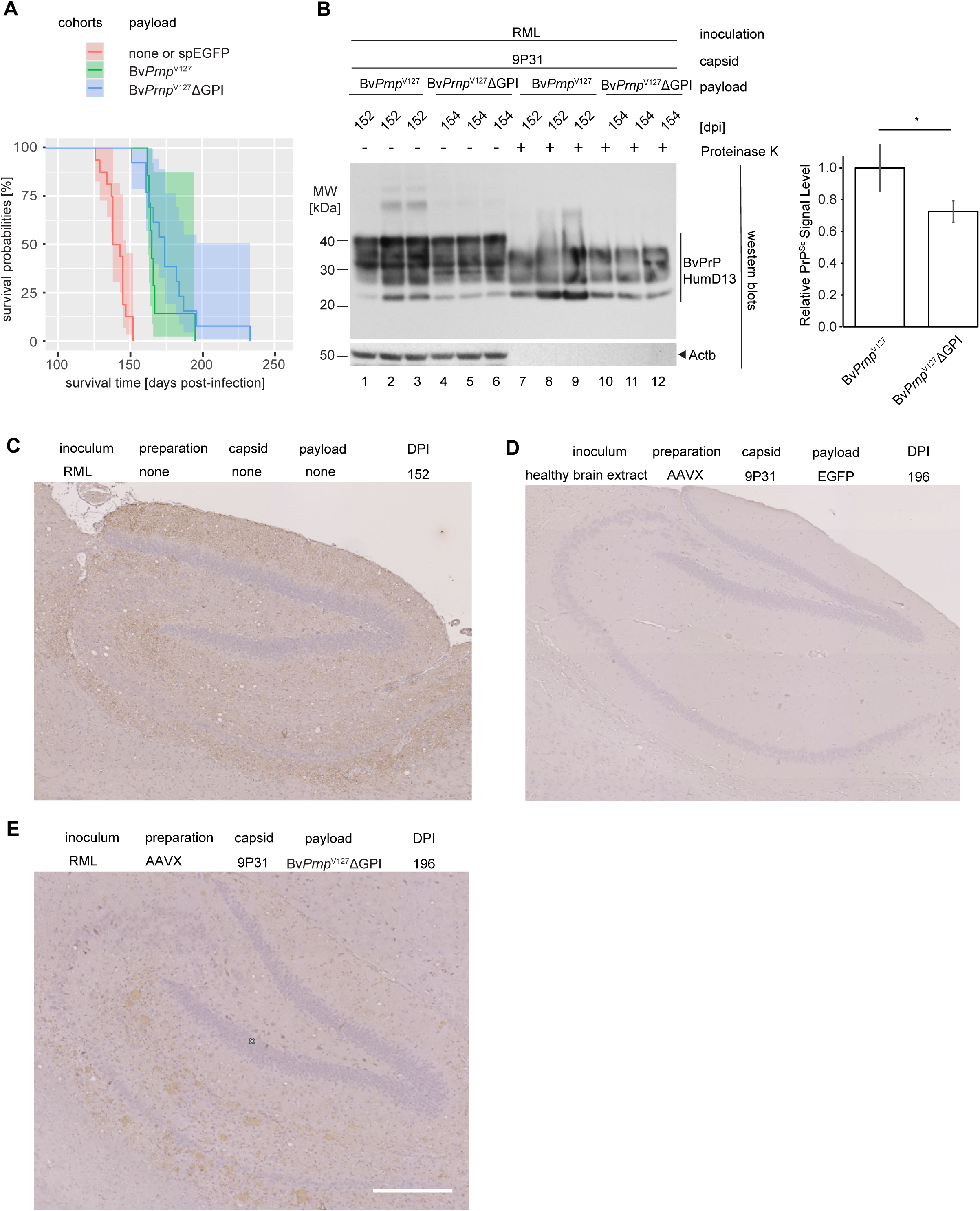
Bv*Prnp*^V127^ΔGPI reduced levels of PK-resistant PrP^Sc^ relative to anchored *BvPrnp^V127^* yet gave rise to a distinct protein deposition phenotype. (A) Comparison of survival extension in cohorts of Bv*Prnp* ki mice following retro-orbital transduction of 9P31 encapsulated virus vectors coding for anchored BvPrP*^V1^*^27^ versus anchorless Bv*Prnp^V127^*ΔGPI. (B) Side-by-side comparison of steady state total PrP levels with and without prior PK digestion in end-stage prion disease Bv*Prnp^V127^*ΔGPI transduced with the respective therapeutic constructs, including quantification of western blot signals. (C) IHC characterization of PrP-reactive signals within the hippocampal formation of an RML-inoculated Bv*Prnp* ki positive control mouse that was not transduced. (D) Negative control IHC image of a hippocampus from a Bv*Prnp* ki mouse that was not prion-inoculated but transduced with a 9P31-EGFP virus. (E) Bv*Prnp* ki mice following retro-orbital transduction of 9P31 encapsulated virus vectors coding for anchorless Bv*Prnp^V127^*ΔGPI at 200 dpi. The IHC images were counterstained with Hematoxylin-Eosin. Sizing bar is 250 µm.

When the brains of 9P31-Bv*Prnp*^V127^ΔGPI-transduced mice were assessed by immunohistochemistry, next to negative control mice that had been RML-inoculated and mock transduced (**Fig 6C**) or non-inoculated mice that were transduced with an EGFP coding vector (**Fig 6D**), the pattern of protein deposition differed. Specifically, a subset of deposits in the hippocampus of mice transduced with Bv*Prnp*^V127^ΔGPI tended to be larger than the corresponding hippocampal PrP deposits in mice transduced with EGFP, which were smaller and more diffuse, as expected for the RML strain (**Fig 6E**). Another report described a predominant perivascular deposition phenotype in transgenic mice engineered to overexpress anchorless mouse *Prnp*ΔGPI without the protective mutation ^11^. No conspicuous association of deposits with blood vessels was observed in 9P31-Bv*Prnp*^V127^ΔGPI-transduced mice but this point warrants further investigation.

### Diagnostic signature of RML prion disease marked by astrocytosis, microgliosis, complement activation, and cellular calcium influx recognizable in asymptomatic mice

To get a better understanding of the biological basis for the survival extension, we next sought to answer the question whether the proteins whose steady-state levels are most increased in prion disease, can also be seen to show the same pattern 50 days ahead of their death in animals that had been transduced with 9P31-Bv*Prnp^V127^*ΔGPI vectors. In designing this experiment, we also considered the potential diagnostic value of proteins whose steady-state levels are altered in the disease, as monitoring their levels cannot only be valuable for early diagnosis but also for gauging treatment success. We focused this analysis on gene products whose steady-state levels are most upregulated, as this would allow them to be positively identified. Consistent with the hierarchical clustering data, this analysis revealed that the treated animals not only showed the same proteins to be upregulated but the levels of their upregulation was mostly consistent, and marked by a relatively high signal-to-noise, i.e., twofold to 25-fold steady-state protein levels relative to naïve mice, suggesting that the proteomic signature of these changes may become recognizable well ahead of the time point chosen for this pilot analysis (**Fig 7**). To understand the cellular context in which the proteins operate whose steady-state levels were most increased, we capitalized on the availability of an in-depth mouse brain single cell transcriptomics dataset ^40^, accessible through the Allen Brain Map, which can be interrogated with mouse gene lists through the ‘Transcriptomics Explorer’ algorithm. When we queried this algorithm with the 50 mouse genes whose protein products were most upregulated in RML-infected mouse brains, it returned almost exclusively assignments to brain cell types that are non-neuronal (**Fig 7, heatmap data**). Several of the expression products of the genes in the list, including GFAP and AIF, are known markers of astrocytosis and microgliosis, respectively, indicating the attempt of RML-infected brains to respond to injury.

**Fig 7.**
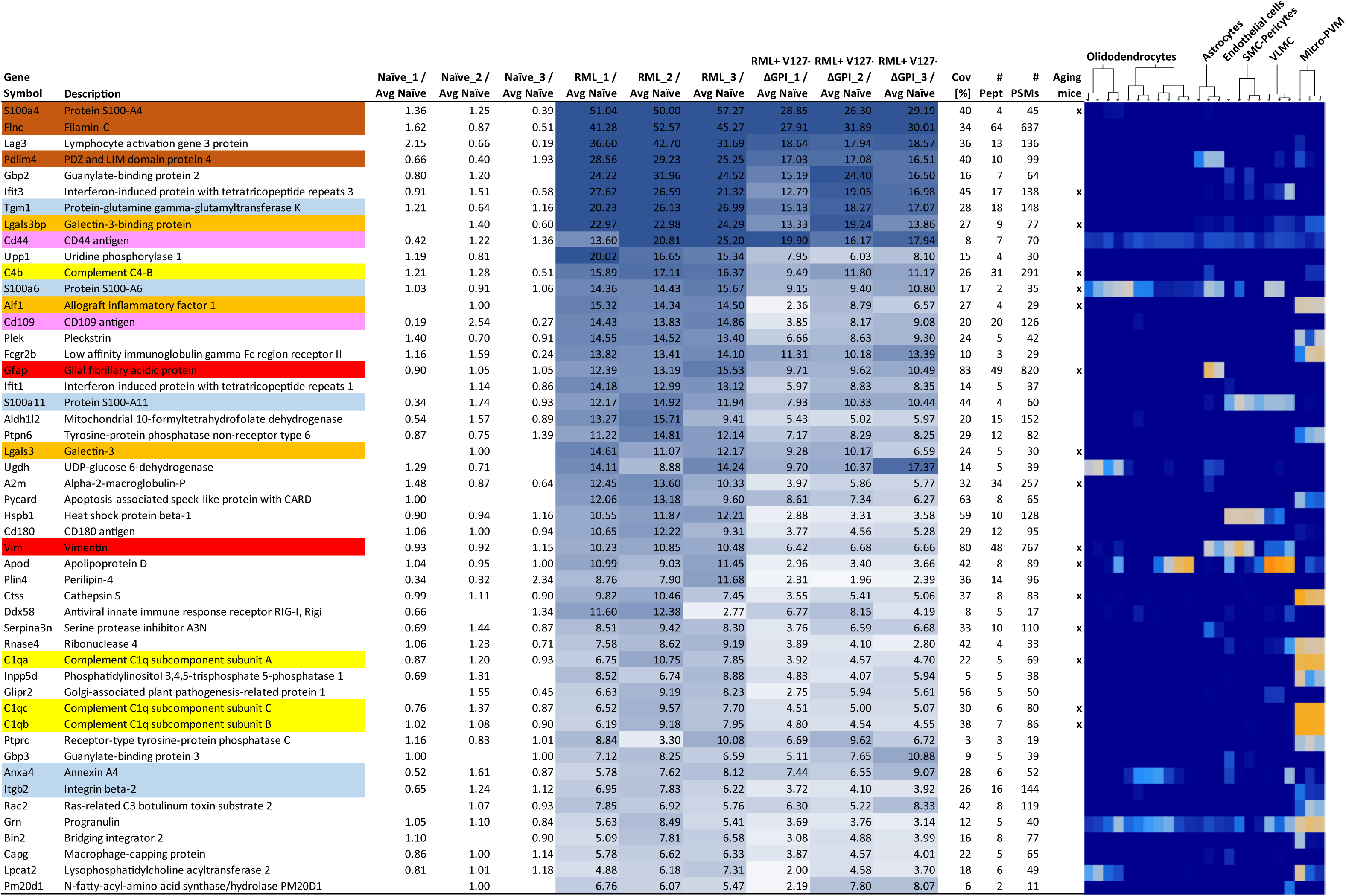
Diagnostic signature of RML prion disease marked by astrocytosis, microgliosis, complement activation, and cellular calcium influx is already recognizable in asymptomatic mice. Shortlist of proteins whose steady-state brain expression levels were most increased in RML-inoculated mice, relative to naïve control mice. Comparing the enrichment levels of individual proteins across the three treatment cohorts reveals that a proteomic signature similar to the 9P31-spEGFP transduced control mice (here shown as RML_1, RML_2, and RML_3) was already present in the 9P31-Bv*Prnp^V127^*ΔGPI transduced mice (here shown as RML+V127-ΔGPI_1, RML+V127-ΔGPI_2, and RML+V127-ΔGPI_3) approximately 50 days before they reached their humane prion disease endpoint. The list is dominated by proteins known to play a role in astrocytosis (highlighted in red color), microgliosis (orange), or the complement response (yellow). Note also the enrichment of proteins known 1) to carry calcium binding domains or to be functionally dependent on calcium (blue colored), 2) to be functionally linked to the reorganization of the actin cytoskeleton (brown), or 3) to form a CD44-CD109 complex (magenta). The column headed ‘Aging mice’, identifies proteins whose transcripts were recently shown to be increased in aging mice ^41^, thereby possibly discouraging their use as specific prion disease biomarkers. On the right, a partial heat map (from the Allen Brain Map) indicates the brain cell types that were previously reported to express most prominently the transcripts of genes shown ^40^.

Since many of the same biological processes are also activated in the ageing brain, we wondered how the shortlist of 50 proteins compares to a similar shortlist of 73 transcripts that were recently observed to be most consistently increased in ageing mice ^41^. Remarkably, 16 of the 50 genes that were most upregulated in prion disease were also most robustly increased in ageing mice. Three additional characteristics of the top 50 gene products are notable and may warrant further investigation to evaluate their usefulness for generating a shortlist of prion disease-specific biomarkers, namely, 1) the list being significantly enriched (*p* = 6.7 E-3) in proteins that carry calcium binding domains or whose function is calcium-dependent (S100a4, S100a11, S100a6, Anxa4, Itgb2, Tgm1), 2) three of the four most upregulated proteins, S100a4, Flnc, and Pdlim4, being functionally linked to actin cytoskeleton organization, 3) the presence of CD44 and CD109 in the list, which are known to form functional interactions and are neither associated with non-neuronal cells nor being activated in aging mice. Taken together, this analysis revealed a signature of prion disease perturbations of potential use for monitoring disease progression that is strongly blunted by the gene therapy and therefore may also serve to gauge treatment success.

### Lower steady-state levels of cell surface proteins residing in proximity to PrP^C^ cannot be accounted for by the loss of neurons that occurs late in the disease

In a prior report, we studied the molecular environment of PrP^C^ in a mouse brain using an *in vivo* crosslinking affinity capture paradigm and had published the top-ranked 40 proteins ^42^. With the deep global proteome analyses at hand, we revisited this shortlist to determine if proteins in proximity to PrP^C^ await a specific fate late in the disease. All but one of the 40 proteins in the original paper were also identified in this global proteome analysis based on at least 10 unique peptides and a range of 185 to 1325 PSMs, thereby affording robust relative quantitations. Interestingly, at late-stage prion disease, none of these PrP^C^ candidate interactors was upregulated, many were approximately two-fold down regulated, and only a few were not affected in their steady-state levels. A common feature of the latter subgroup seems to be that they encounter the prion protein within its passage along the secretory pathway, whereas the downregulated proteins reside next to PrP at the cell surface. Many of these proteins belong to the subset of proteins that were most strongly and consistently downregulated in the entire dataset (**Fig 8A**). In fact, if proteins which we had reported to reside in proximity to PrP at the cell surface were captured in a KEGG pathway, its downregulation would emerge among all known KEGG pathways annotations as a highly significantly downregulated pathway (*p* = 2.37E-25), with only three cell surface proteins, namely contactin-1, the amyloid precursor protein, and the myelin associated glycoprotein, bucking this trend.

**Fig 8.**
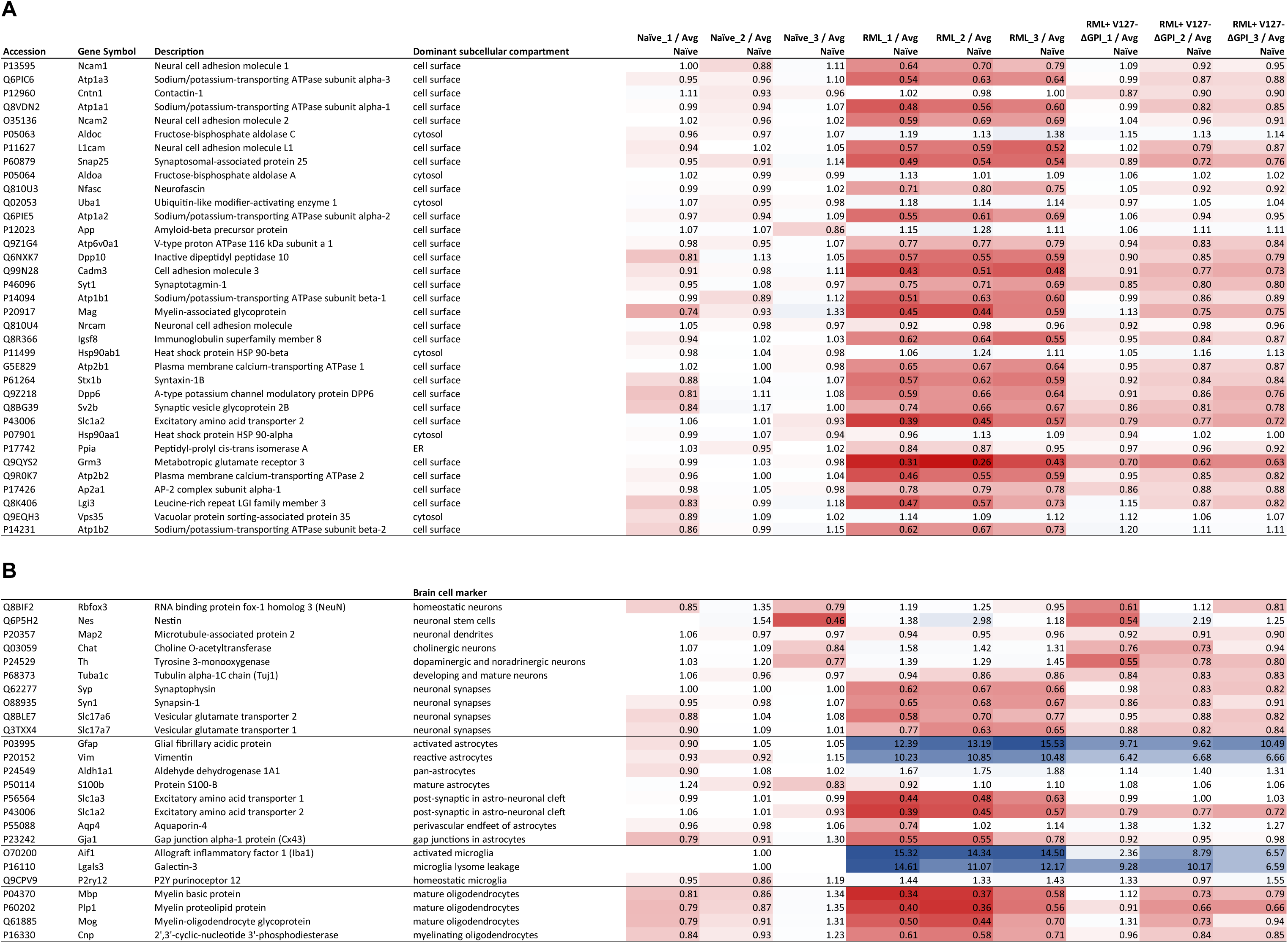
Lower steady-state levels of cell surface proteins residing in proximity to PrP^C^ cannot be accounted for by the loss of neurons that occurs late in the disease. (A) Proteins reported to reside in proximity to PrP^C^ at the cell surface are downregulated in end-stage RML prion-inoculated mice. The list depicts previously reported proteins in an affinity-capture PrP interactome from an *in vivo* crosslinked mouse brains ^42^. The intensity of the red shading reflects the level of reduction of a given protein entry relative to its average steady-state levels in age-matched naïve control mice. Note the consistent red shading in proteins whose dominant subcellular compartment is the cell surface. (B) Although neurons are understood to die in late-stage prion disease, the steady-state levels of several neuronal markers (NeuN, nestin, or Tuj1) are not noticeably lowered in a sagittal half brain extracts generated at the prion disease end-stage, indicating that for a majority of brain neurons the disease merely causes the retraction of synapses but not their full disappearance. The table also shows well-known brain cell marker proteins that are preferentially expressed in astrocytes, microglia, or oligodendrocytes, indicating the activation of astrocytosis and microgliosis (recognizable by the blue shading that reflects upregulation), as well as consistent loss oligodendrocytes (recognizable by the consistent red shading of all oligodendrocyte marker proteins).

We next considered if a trivial explanation of neurons dying in late-stage prion diseases might account for this selective downregulation of cell surface proteins in proximity to PrP, since most of them are associated with neurons. To address this question, we extracted from the global proteome dataset the relative quantities of proteins, which are commonly associated with specific brain cell types. Specifically, this incomplete list was comprised of ten neuronal, seven astrocytic, four oligodendrocytic, and two microglia markers (**Fig 8B**). Within the ten neuronal markers, we observed that steady-state level of NeuN and Tuj1, proteins that are frequently used to quantify homeostatic and mature neurons, did not change in late-stage-disease. Neither did other neuronal markers which identify neuronal stem cells (Nestin) or specific subtypes of neurons (choline-O-acetyltransferase for cholinergic neurons and tyrosine 3-monooxygenase for dopaminergic or noradrenergic neurons). Only neuronal markers associated with synapses, including Synaptophysin or Synapsin, were observed to be downregulated. These results indicate that the mere neuronal death that occurs late in the disease does not account for the reduction in the steady-state levels of the previously reported interactors and candidate interactors of the prion protein.

## DISCUSSION

This report described results from a proof-of-concept study, which evaluated the therapeutic potential of a virus-delivered gene therapy based on a secreted bank vole PrP^V127^ΔGPI expression product. The study documented an approximately 50-day survival extension in RML prion-inoculated mice using this approach. The subsequent analyses of brain samples indicated that this survival extension was obtained irrespective of whether the heterologous therapeutic protein was expressed at low or high levels, suggesting a ceiling of therapeutic potency in this paradigm. A deep proteomic analysis of brain samples, collected at the time when negative control mice succumbed to prion disease, yet the treated mice were still free of overt symptoms, showed that the heterologous expression of Bv*Prnp*^V127^ΔGPI mitigated proteomic perturbations observed in non-treated RML-inoculated mice and was not in itself causing specific changes to the proteome. Mice that were transduced with rAAV vectors coding for anchored Bv*Prnp*^V127^ led to a shorter average survival extension of 25 to 30 days, consistent with the interpretation that cross-correction contributed to the enhanced potency of the secreted construct. Further investigation of the global proteome dataset established that both prion disease and Bv*Prnp*^V127^ΔGPI overexpression (albeit it in a delayed manner) were associated with astrocytosis and microgliosis, as well as a deterioration of synapses and proteins underpinning the sleep-wake cycle. The proteomic data also revealed that the disease causes brain cells to invest in a rescue biology centered on cellular Ca^2+^ influx and a replenishment of cell surface proteins, prominently manifest in increases in steady state levels of an endoplasmic reticulum protein processing subproteome and components of the spliceosome.

Since prion diseases require templated polymerization for their propagation, any mismatch in the prion sequence has the potential to hinder disease progression. The idea to harness the power of sequence variants for therapy is not new. In the prion field, this idea has had traction ever since sequence variants that manifest in animal and human populations as polymorphisms or mutations were shown to confer partial or full protection against prion diseases. Initial work in this area was mostly based on the transient or stable transfection of protective PrP mutants *in vitro* based on the ScN2a cell model and the measurement of PK resistance as a surrogate for disease burden ^43, 44^.

To date, *in vivo* work investigating this concept has remained limited. For instance, it was shown that expressing the protective human prion gene polymorphisms Q167R or Q218K at the same level as wild-type PrP could slow prion disease in mice after prion inoculation but did not prevent it completely, unless the protective transgene was exclusively expressed ^45^. A follow-on paper by one of the authors documented that the protective K218 variant could reduce the burden of PK-resistant PrP^Sc^ in ScN2a cells when the protein was added to the cell culture medium in recombinant form, lacking N-glycans and GPI anchor ^46^. The subsequent infusion of the same recombinant protein into mouse brains through an intracerebroventricular catheter was reported to have prolonged the prion disease incubation period from 117 days to 131 days ^47^. A separate report based on the same therapeutic concept, but a different means of delivery documented that the lentiviral transduction of a gene therapy coding for the protective R167 variant achieved a 30-day survival extension in prion-inoculated mice ^48^. Several experimental differences to the work reported here stand in the way of interpreting differences in outcomes, including that the protective R167 mutant was expressed in the context of a mouse sequence that coded for the attachment of a GPI anchor, the use of the Me7 prion inoculum, the administration of the treatment through an intracerebral cannula implant, and the choice to administer repeat treatments at 80 and 95 dpi.

To our knowledge, this report is the first to test the protective effect of the V127 mutation *in vivo* using a paradigm that introduced the therapeutic protein through viral delivery. Key differences of our experimental design to the natural protective V127 heterozygosity that evolved in the Kuru endemic region in Papua New Guinea are:

1. The heterologous germ-line expression of this protective mutant in all cells in humans reported to carry this mutation naturally versus the heterologous expression following the transduction of a subset of brain cells weeks after the brain had been exposed to the prion agent.
2. The human sequence context of the naturally evolved V127 mutation versus our decision to work with the BvPrP sequence. We had previously shown that the V127 mutation retains at least some of its protective capacity when it is embedded in the BvPrP sequence by showing that mouse CAD5 cells, in which we abrogated the expression of the endogenous mouse *Prnp* gene and instead expressed Bv*Prnp*^V127^ as a transgene, became resistant to prion infection ^49^. As we prepared this manuscript for publication, this finding was expanded on by another team; using the same CAD5 cell paradigm, as well as N2a cells, the authors showed that the protection conferred by V127 (V126 in mice) extends to several natural and artificial prion strains ^50^. In a separate branch of their work the same team showed that the lentiviral delivery of the V127 mutation to primary cultures of hippocampal neuronal from PrP-null mice can prevent the prion-induced retraction of dendritic spines ^51^.
3. The expression of the naturally evolved protective PrP^V127^ mutant as a GPI-anchored protein versus our choice to secrete BvPrP^V127^ΔGPI to achieve cross-correction (**Fig 9**).

**Fig 9.**
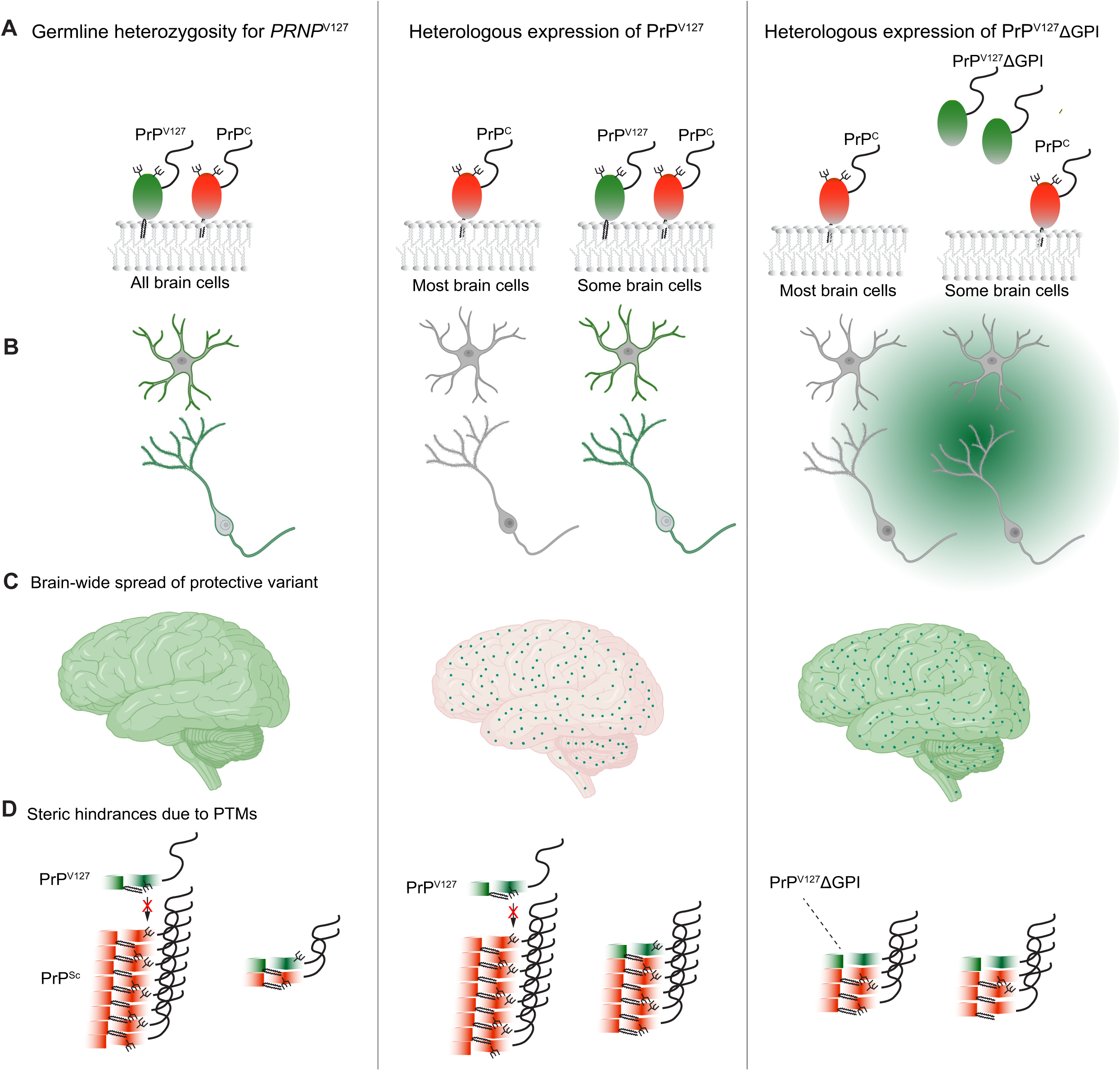
Model interpreting results and highlighting facets of ECC that may translate into advantages of expressing PrP^V127^ΔGPI over anchored PrP^V127^. (A) Whereas in germline-expressing heterozygous kuru survivors carrying the V127 mutation, PrP^V127^ is expressed in all cells, only a subset of cells will be transduced when the protective mutant is delivered through rAAV vectors. This deficiency may be partially compensated through an increase in the half-life of PrP^V127^ΔGPI, relative to PrP^V127^. (B) The secretion of PrP^V127^ΔGPI promotes local spread, thereby promoting access to extracellular PrP^Sc^ seeds. (C) Whereas the germline expressed PrP^V127^ is present in all brain areas due to its ubiquitous expression (symbolized by the smooth green color), PrP^V127^ΔGPI can be highly expressed only in a subset of brain cells. However, a strong promoter may allow it to achieve a similar brain-wide protection through its diffusion within extracellular spaces and the CSF. (D) The presence of a GPI-anchor and N-glycans has the capacity to slow access to nascent PrP^Sc^. This may not be a hindrance if PrP^V127^ is germline expressed alongside wild-type PrP as the immediacy of contact between the conversion susceptible and refractory isoforms may still provide the earliest possible protection (here depicted as a capping mechanism, one of several possible scenarios through which the V127 may protect in a dominant negative fashion). Note that the size of PrP^Sc^ seeds is meant to indicate the readiness with which the V127 mutant can block aggregation, i.e., a capped aggregate comprising just one PrP^Sc^ and one V127 mutant molecule is meant to indicate ready inhibition, whereas four and eight building blocks in a PrP^Sc^ aggregate symbolize moderate and impaired inhibition of PrP^Sc^ accumulation, respectively. To simplify the cartoon, only one N-glycan is shown, and steric hindrances are indicated with red-crossed arrows.

Some level of cross-correction may occur naturally, because a subset of PrP^C^ and its PrP^V127^ mutant are shed by a-disintegrin-and-metalloproteinase 10 (ADAM10) from the cell surface through endoproteolytic cleavage at a site located a few residues upstream of the GPI-attachment site ^52^, thereby giving rise to endoproteolytic products referred to as shed PrP^C^ (sPrP^C^ and sPrP^V127^) ^53^. However, the natural proportion of sPrP^C^ to total PrP^C^ is low, estimated to be 7-10% in rodents ^54^. Therefore, the natural ADAM10-mediated release of PrP^C^ would provide limited protection to the brain if only a small subset of cells can be made to express the GPI-anchored PrP^V127^ mutant.

As far as we are aware, this report is also the first to capture changes to the levels of almost 5,000 proteins in late-stage prion disease. The value of this dataset exceeds its limited use in this study. For instance, the knowledge of proteins whose levels are profoundly up- or down-regulated may lead to disease-specific diagnostic markers that can be used to track disease progression. Although a few proteins, including GFAP ^55^, have long been known to be altered in their steady-state levels in prion disease, these proteins may have limited value as disease-specific markers and, as we showed, are also not the proteins whose levels are most altered in the disease. Neuronal cell surface proteins whose levels increase in a manner that tracks with disease progression, may not only be of diagnostic value but may also represent attractive targets for guiding therapies to the specific neurons that are most affected in the disease.

PrP^C^ is thought to play also a central role as a mediator of cellular toxicity in Alzheimer’s disease (AD), where it has been shown to act as the cell surface receptor necessary for small oligomeric assemblies of the Abeta peptide (oAβ) that are endoproteolytically released from the larger amyloid precursor protein (APP) ^56^. Consistent with this scenario, reduced levels of cell surface PrP^C^ have been reported to prevent toxicity in AD models ^56–63^. Although not formally tested in this context, the small structural differences reported for PrP^V127^ cannot be expected to interfere with its ability to serve as a receptor for oAβ since the binding site maps to the disordered N-terminal domain of PrP that is shared between wild-type PrP and PrP^V127^ ^56^. The secretion of PrP^V127^ΔGPI would be expected to ameliorate the risk of toxicity in AD by competing with cell surface PrP^C^ for binding of toxicity-inducing entities and, consequently, sequestering them away from the cells.

This proof-of-concept study had limitations. Most striking amongst them are the limited survival extension that was observed in the cohort of mice transduced with virus particles coding for PrP^V127^ΔGPI despite the pronounced brain-wide expression of the protective construct achieved. We are considering several potential causes for this limitation. We wonder if the BvPrP sequence, notorious for its standout propensity to convert easily to BvPrP^Sc^ and to act as a universal acceptor for prion strains from a wide range of species, might have blunted the protective capacity of the V127 mutation. A related concern is that little is known about how well dominant negative mutations translate across species. There are many data points which established that several human PrP polymorphisms and mutations can confer protection also when inserted into the prion sequences of other species, most notably the mouse *Prnp* sequence. However, it would be surprising if all dominant negative mutations translated equally well in such cross-species experiments. In fact, unpublished work by others appears to indicate limits in the protection that the expression of V127 can confer toward certain prion strains, including RML. Specifically, a preliminary presentation of transgenic mice expressing mouse *Prnp*^V126^ in their germ-line revealed that these mice succumbed to RML prion disease albeit after a delayed disease course ^64^. Thus, with the wisdom of hindsight it seems that our choice to work with the RML prion strain for this pilot study was poorly suited to assess the protective potency of a PrP^V127^ΔGPI therapy *in vivo*. Future work based on mice expressing human *PRNP* as a knock-in ORF, inoculated with human prion inocula, and treated with a virus-delivered gene therapy based on the expression of human PrP^V127^-ΔGPI may reveal the true impact of this limitation.

A weaker argument can be made that the treatment was administered too late. It has repeatedly been described that it takes approximately three weeks for transduced rAAV vectors to unleash the full expression of their payload ^65^. Our choice to administer the treatment after 60 days therefore may have led therapeutic PrP^V127^ΔGPI levels to plateau only after approximately 81 dpi. When viewed considering observations from the ASO treatment studies where a marked decline in therapeutic potency was observed when the therapy was administered later than 78 dpi, one may be tempted to conclude that our therapy came too late to confer its full protective effect ^4^. However, this interpretation is tempered by a recent report on rAAV-delivered ZFRs, which documented that pronounced survival extension occurred even when this treatment modality was administered at 120 dpi ^7^. Finally, a potential concern is that we observed subtle changes to the protein deposition phenotype in RML-infected mice when the ratio of protective BvPrP^V127^ΔGPI to wild-type expression was 3 : 1. This concern may be less worrisome when considering that this expression level was unnecessarily high.

## CONCLUSIONS

Results from this pilot study may be viewed both as disappointing and hopeful. Disappointing due to the limited survival extension of 50 days achieved. The data are hopeful because they validate the therapeutic concept of a rAAV-delivered gene therapy based on the V127 protective mutation. Moreover, the data substantiate our hypothesis that cross-correction afforded by the secretion of the anchorless protective mutant can supersede the protection offered when the V127 mutation is expressed in the context of GPI-anchored PrP. Finally, we were encouraged to find that the retro-orbital intravenous delivery of a brain penetrant gene therapy can lead to steady-state PrP^V127^ΔGPI levels that exceeded 3 : 1 the endogenous expression levels of this highly expressed endogenous protein. This is hopeful as mice had been shown to be protected against human prion inocula when their germline encoded PRNP^V127^ and PRNP genes were present at a ratio of 1 : 3 ^9^, i.e., we may have considerable titration room left to address this aspect of the gene therapy.

We also remain hopeful that the survival extension ceiling encountered in the Bv*Prnp* sequence context can be lifted by moving this approach to human prion gene sequences and human prion inocula. In the context of human familial disease, it will be of interest to learn if it is favorable to deliver the V127 protection within a bespoke therapeutic construct that also comprises the specific disease-causing inherited mutation of the individual treated. It seems plausible that this additional design element would favor the disease blocking interactions of the protective protein with the respective disease-causing PrP^Sc^ conformers.

## MATERIALS AND METHODS

### Antibodies

Primary antibodies: Human monoclonal anti-prion F(ab’)2 antibody (epitope: residues 96-104 in mouse PrP) ^66^, clone HuM-D13, 1:5,000, generously provided by the laboratory of Dr. Emil F. Pai (University of Toronto, ON, Canada). Rabbit monoclonal anti-prion IgG antibody (epitope: 221SQA223 and Y225), clone EP1802Y, 1:10,000 (catalog number ab52604, Abcam, Cambridge, United Kingdom). Mouse monoclonal anti-beta-actin (Actb) IgG2b antibody, clone BA3R, 1:40,000 (catalog number MA5-15739-HRP, Thermo Fisher Scientific, Waltham, MA, USA).

Secondary antibodies: Goat anti-human IgG F(ab’)2 horseradish peroxidase (HRP) conjugated antibody, 1:5,000 (catalog number 31414, Thermo Fisher Scientific). Goat polyclonal anti-rabbit HRP secondary IgG antibody, 1:5,000 (catalog number 31460, Thermo Fisher Scientific).

### Cloning

The self-complementary therapeutic Bv*Prnp* vectors were built using Gibson assembly. Briefly, the synthetic Bv*Prnp*^V127^ sequence avoiding CpG motifs was purchased from a local gene synthesis service (Bio Basic Inc, Markham, ON, Canada). The backbone of the recombinant AAV transfer plasmid with its regulatory elements, designated as pscAAV-CBh-Null-WPRE3-enSV40pA, were a gift from the Michael J Fox Foundation (catalog number 194245, Addgene, Watertown, MA, USA). The transfer plasmid was opened with StuI and AgeI restriction enzymes. A parallel Gibson assembly, which removed the GPI signal sequence and inserted a canonical Kozak sequence and translation termination nonsense codon, was undertaken with a PCR reaction mix composed of Q5 High-Fidelity 2x Master Mix (catalog number M0492S, New England Biolabs, Ipswitch, MA, USA), template DNA and the following primers:

Forward: 5’– tcaggttggaccggctagcaccggtgccaccATGGCCAACCTcagctactg – 3’,

Reverse: 5’ – gtaatccagaggttgattaggTCATCTGCCCTCATAGTAGGCCTGGG – 3’.

The reaction ran for 10 cycles with an annealing temperature of 55 °C, followed by 20 cycles at the elevated annealing temperature of 65 °C. Finally, the gel-purified open vector backbone and Bv*Prnp*^V127^ or Bv*Prnp*^V127^ΔGPI sequences were assembled with the help of the HiFi DNA Assembly Master Mix (catalog number E2621L, New England Biolabs) during a 1-hour incubation at 50°C, then transformed into NEB Stable Competent *E. coli* (catalog number C3040H, New England Biolabs).

### Cell culture and rAAV vector production

HEK293T cells were maintained in DMEM (catalog number 119650-92, Thermo Fisher Scientific) supplemented with 50 U/mL Pen/Strep (catalog number 15140122, Thermo Fisher Scientific), NEAA (catalog number 11140-050, Thermo Fisher Scientific) and 10% FBS (catalog number 12483020, Thermo Fisher Scientific). Cultures were grown at 37°C, in an atmosphere of 5% CO_2_ and 95% relative humidity. The media were replaced every 48 to 72 hours and the cells were passaged at 90% confluency in using 0.25% Trypsin-EDTA (catalog number 15050065, Thermo Fisher Scientific). The rAAV vector production and purification followed a previously established protocol ^34^. In brief, HEK293T cells were cultured in a HYPERFlask (1,720 cm² surface area) with 560 mL of growth medium (catalog number CLS10031-4EA, MilliporeSigma). Upon reaching 70-80% confluence, the cells were transfected with three plasmids: an AAV packaging plasmid containing Rep and Cap genes, a helper plasmid encoding adenovirus genes E2A, E4, and VA, and the rAAV transfer plasmid. Four days post-transfection, the cell culture medium was supplemented with 3 mL of Triton-X 100 (catalog number ×100, Sigma-Aldrich), 250 µl of RNAse A (catalog number EN0531, Thermo Fisher Scientific), 56 µL of Pluronic F-68 (catalog number 24040-032, Thermo Fisher Scientific), and 56 µL of Turbonuclease (catalog number T4330, Sigma-Aldrich). The mixture was agitated at 150 RPM and 37 °C for 1 hour. Afterward, the lysate was collected and the HYPERFlask was washed with 140 mL of Dulbecco’s phosphate-buffered saline (PBS) (catalog number 14190-144, Thermo Fisher Scientific). The combined lysate and PBS wash were then centrifuged at 4,000 × g (RCF) for 30 minutes to pellet the cell debris. Finally, the supernatant was filtered through a 0.45 µm PES Autofil bottle top filter (catalog number 1143-RLS, Foxx Life Sciences).

### rAAV vector purification method

The purification of rAAV vectors made use of 50 µm POROS CaptureSelect AAVX resins (catalog number A36652, Thermo Fisher Scientific). The resin was washed with 20 column volumes (CVs) of Low Salt Wash Buffer (50 mM Tris/HCl, pH 7.4, 150 mM NaCl, 0.01% Pluronic™F-68 surfactant, 1% Triton-X100). The affinity capture step was initiated by the slow (0.3 mL/min) loading of rAAV vectors harvested from HEK293 supernatants onto the AAVX resins. Once the rAAV vectors were loaded, the resin was washed with 20 CVs of Low Salt Wash Buffer, 20 CVs of High Salt Wash Buffer (300 mM NaCl, 50 mM Tris/HCl, pH 7.4, 0.01% Pluronic™F-68 surfactant), and 20 CVs of Low Salt Wash Buffer without Triton-X100. Elution was induced by pH drop (0.2 M glycine and 0.01% Pluronic F-68, pH 2-2.5) and led to the collection of 4 mL fractions, which were rapidly pH neutralized with 420 µL of neutralization buffer (1 M Tris/HCl, pH 8, 0.1% Pluronic F-68). Prior to use, rAAV preparations were filtered through 0.22 µM PES membranes using a Stericup Quick Release-GP filtration system (catalog number S2GPU11RE, MilliporeSigma Canada Ltd., Oakville, ON, Canada).

### qPCR

Quantitative PCRs were undertaken to determine virus titers using a well-developed protocol (Challis et. al. 2019). Briefly, serial dilutions of DNA standards were generated using the respective transgene plasmids which were linearized with ScaI (catalog number R3122S, New England Biolabs). Viral preparations were digested with 50 U/mL of DNase I (catalog number EN0521, Thermo Fisher Scientific) in 2 mM CaCl^2^, 10 mM Tris-HCl, 10 mM MgCl^2^. Next, viral capsids were digested using Proteinase K (catalog number EO0491, Thermo Fisher Scientific) in 1M NaCl, 34 mM N-lauroylsarcosine. The final qPCR measurement was conducted using a LightCylcler 480 II (catalog number 05015278001, Roche Diagnostics, Indianapolis, IN, USA) with the help of the SYBR green master mix (catalog number 4367659, Thermo Fisher Scientific) as well as the following primers which target the CBh promoter: Forward: GTTACTCCCACAGGTGAGC and reverse: CCAACCAACCATCCCTTAAAC.

### Animals

Animal studies were necessary to assess the *in vivo* efficacy of a rAAV vector delivered gene therapy that capitalizes on the protective V127 variant. All animal procedures were based on guidelines by the Canadian Council on Animal Care and were authorized by the University Health Network (UHN) Animal Care Committee (Animal Use Protocol 6840). All personnel involved in animal care or surgical procedures received specialized training for their respective tasks to ensure the humane treatment of the animals. C57BL/6J mice were obtained from the Princess Margaret Cancer Research Centre (University Health Network, Toronto, ON, Canada). Homozygous Bv*Prnp* I109 ki mice (Background: C57BL/6J) were described before ^67^. A maximum of five mice per cage were kept at artificial 12-hour day and night cycles, drinking water *ad libitum*, and given 18% protein chow as solid food source. The mice received daily checks to assess their health and appearance. Their cages were changed weekly. To immobilize mice to be inoculated, transduced with rAAV vectors, or euthanized, anesthesia was induced with inhaled 5% isoflurane. Subsequently, when survival of the animals was intended, isofluorane levels were maintained at 2%. The anesthetic depth was assessed by performing a toe pinch. Respiration was observed during surgery to ensure a regular respiratory pattern. Prion-inoculated mice were closely monitored for signs of distress or pain, and humane endpoints were predefined (>20% weight loss, abnormal posture, lethargy, and impaired ambulation). Mice meeting these criteria were euthanized. Nesting scores were assigned based on whether the mice built no nest (score = 0), built a flat nest (score = 1), or built a three dimensional nest (score = 2). The nesting material consisted of pulped cotton fiber supplied in sheets that break easily into nestlets (catalog number NES3600, Ancare Corp., Bellmore, NY, USA).

### Intracerebral prion inoculations

Intracerebral prion inoculations were undertaken with 4-6-week-old mice. The inoculant was generated by infecting a C57BL/6J mouse with RML-prions, then sacrificing the animal when terminally ill with prion disease and homogenizing the brain to a final concentration of 3% brain extract in PBS (v/v). Following deep anesthesia, mice were free-hand injected into the right parietal lobe 20 µL of inoculant at a depth of 3 mm with a 29-gauge SafetyGlide insulin syringe (catalog number 305930, Becton Dickinson Canada, Mississauga, ON, Canada).

### Retro-orbital injections

Ahead of this procedure, the mice were anesthetized as described in the ‘Animals’ section. Additionally, all mice received for this procedure one eye drop of 0.5% proparacaine hydrochloride ophthalmic solution as a local anesthetic before injection. The injections were administered into the right orbital sinus using a 28-gauge Micro-Fine IV insulin syringe (catalog number 329420, Becton Dickinson Canada). All animals received a single retro-orbital injection of 1×10¹² viral genomes (vg), diluted in PBS to a final volume of 100 µL. Six-week old female C57BL/6J mice were injected with 9P31-spEGFP to test the CNS spread after retroorbital injection.

### Mouse tissue collection

Mice were deeply anesthetized with isoflurane and euthanized by six-minute transcardiac perfusion with PBS. Next, the brains were carefully extracted, their meninges removed, and their brains split into two sagittal hemispheres. The right hemisphere was designated for biochemical analyses and kept at −80°C until homogenization. The left hemisphere was post-fixed for subsequent pathological analyses by immersion in 10 mL of neutral-buffered formalin (catalog number HT501128-4L, Sigma-Aldrich). If mouse hemispheres were dedicated assigned for cryo-sectioning, the post-fixation was limited to two hours at 4°C.

### Cryo-sectioning of mouse brains for direct fluorescence detection

Mouse brains for cryo-sectioning were dissected at three weeks post-injection. After fixation, brains were cryoprotected by immersion in 30% sucrose in PBS at 4°C for up to 36 hours causing the brain tissue to sink. Brains were then incubated in a 1:1 mixture of 30% sucrose and Tissue-Tek O.C.T. Compound (catalog 25608-930, VWR, Radnor, PA, USA). Next, brains were positioned in a cryomold (catalog number 70182, Electron Microscopy Sciences, Hatfield, PA, USA) and embedded in O.C.T for another one hour at 4°C degrees. Finally, the brains were frozen by partially immersing the cryomolds in a liquid nitrogen-chilled 2-methylbutane bath for 2-3 mins until the O.C.T completely froze. The frozen blocks were briefly air-dried on dry ice to remove residual 2-methylbutane and stored at −80°C.

Before cryo-sectioning, brains were allowed to warm up in the Cryostat chamber (model HM525 NX, Thermo Fisher Scientific) to a temperature of −21°C during a two-hour acclimatization period. Finally, 16 µm sagittal sections were cut and collected using Kawamoto’s film method on a piece of adhesive Cryofilm (catalog number C-FUF303, Section-Lab Co. Ltd., Yokohama, Kanagawa Prefecture, Japan) ^68^. Before imaging, sections were washed for five minutes in PBS and mounted with mounting media containing DAPI (catalog number ab104139, Abcam).

### Immunohistochemical analyses of mouse brain sections

To prepare samples for immunohistochemical analyses, mouse brains were immersed for one hour in formic acid, then rinsed with water, and placed in a Leica Pearl for processing. After sectioning, the slides were dried overnight at 37°C. Ahead of the immunohistochemical staining, the slides were baked for an hour at 60°C to help the tissues to adhere better to the slides. Next the slides were dehydrated through a xylene and ethanol series. The immunodetection made use of the Mouse on Mouse (M.O.M) Elite Immunodetection Kit (catalog number PK-2200, Vector Laboratories, Newark, CA, USA) using the manufacturers instructions. As part of these instructions, the tissue sections underwent antigen retrieval with citrate, pH 6.0, and heat using a commercial Antigen Decloaker formulation (catalog number CB910M, Biocare Medical, Pacheco, CA, USA). Brain slices were incubated with a monoclonal mouse PrP antibody (clone 9A2) at a 1:500 dilution (Wageningen Bioveterinary Research, Lelystad, Netherlands) for 30 min at 22 °C and images developed using the NovaRed system (Vector Laboratories Inc., Newark, CA, USA). Finally, brain tissue sections were counter-stained with hematoxylin and eosin, rapidly dehydrated, and cover-slipped for microscopy analyses.

### Microscopy

Brain samples were imaged under #1.5 glass coverslips (catalog number 48393-060, VWR) on Fisherbrand Superfrost Plus microscope slides (catalog number 22-037-246, Thermo Fisher Scientific) using a Zeiss AXIO Observer 7 inverted LED fluorescence microscope (Carl Zeiss Canada Ltd., North York, ON, Canada). To reconstruct full sagittal brain views, individual fields of view were acquired sequentially across each section and digitally stitched into a composite image using ZEN Blue microscopy software (Carl Zeiss Canada Ltd.).

### Homogenization and protein extraction

Mouse brain tissue was homogenized in 100 mM Tris-HCl (pH 8.3) using a Minilys homogenizer (catalog number P000673-MLYS0-A, Bertin Technologies, Rockville, MD, USA) and Zirconia beads (catalog number 11079110zx, Biospec, Bartlesville, OK, USA) with three sets of 30 second bead-beating pulses. The resulting 20% homogenates were aliquoted, and a protease inhibitor cocktail (catalog number 78425, Thermo Fisher Scientific) was added to all samples that were not designated for subsequent Proteinase K digestion. To extract proteins, a detergent stock solution giving rise to final concentrations of 0.5% deoxycholic acid (DOC) (catalog number DCA333.50, BioShop, Burlington, ON, Canada) and 0.5% NP-40 (catalog number NON505.100, BioShop) was added to the homogenate, followed by gentle vortexing and 30 minute incubation on ice. Next, insoluble debris were removed during consecutive five-minute spins at 500 × g and ten-minute spins at 5,000 × g (RCF). Total protein concentrations of the samples were determined using the Pierce BCA Protein Assay Kit (catalog number 23225, Thermo Fisher Scientific) and subsequently adjusted.

### gDNA extraction and restriction digest

The gDNAs of study mice were extracted from brain homogenates with the Monarch Spin gDNA Extraction Kit (catalog number T3010S, New England Biolabs). For the subsequent PCR reaction, 50 ng of gDNA was amplified with Q5 Hot Start High-Fidelity 2X Master Mix (catalog number M0494S, New England Biolabs) and forward and reverse primers (final concentration 500 nM) that map to gDNA sequences shared by the endogenous and heterologous *BvPrnp* ORFs. Equal volumes of PCR product were then subject to a restriction enzyme (RE) digest with 1 µL of EagI (catalog number R3505S, New England Biolabs), CsiI (catalog number FD2114, Thermo Fisher Scientific), or both REs for cutting the endogenous or the heterologous amplicons or both, respectively. The RE digestion products were finally separated on a 1% agarose gel containing SYBR safe DNA gel stain (catalog number S33102, Thermo Fisher Scientific). The fluorescent image of the gel was captured with the ChemiDoc XRS+ System (Bio-Rad Laboratories, Hercules, CA, USA). Forward: AAGAAGCGGCCAAAG and reverse: TAGTAGGCCTGGGACTC.

### PNGase F digest

To remove N-linked glycans by digestion with PNGase F (catalog number P0704S, New England Biolabs, Ipswich. MA, USA), we made use of supplier provided buffers. Specifically, equal amounts of proteins were denatured using 10× Denaturing Buffer at 95°C for 10 minutes. Next, NP-40, GlycoBuffer, and PNGase F enzyme were added to the denatured samples. To inhibit undesired proteolytic activity, 3 mM phenylmethanesulfonyl fluoride (PMSF) (catalog number PMS123.5, BioShop) was added to the reaction tubes. Next, samples were incubated for 2 h at 37°C with gentle shaking. Finally, an equal volume of 2× LDS sample buffer containing 5% β-mercaptoethanol (BME) was added directly to the samples to stop the reaction and prepare the samples for western blot analyses.

### Proteinase K digest

To assess the relative PrP^Sc^ content of brain extracts, 200 µg each of BCA-adjusted total proteins were treated with Proteinase K (catalog number 25530049, Thermo Fisher Scientific) at a final concentration of 50 µg/mL and a ratio of total protein to Proteinase K of 1:20. Next, samples were incubated for 45 min at 37°C under gentle shaking. The digestion was terminated by adding PMSF to a final concentration of 2 mM, followed by the addition of 2% Sarkosyl (catalog number SLS002.100, BioShop). Next, the digests were ultracentrifuged for 1 hour at 48,000 RPM and 4°C using an Optima TLX Ultracentrifuge (Beckman Coulter, Brea, CA, USA). Finally, the pelleted insoluble PrP^Sc^ was resuspended in 1x Bolt LDS sample buffer (catalog number B0007, Thermo Fisher Scientific), heat-denatured at 95°C for 10 minutes, then analyzed by western blotting.

### Western blotting

To detect total PrP within brain extract samples, proteins were denatured in 1x Bolt LDS sample buffer at a final concentration of 2 µg/µL. Next, samples were heated at 95 °C for 10 min, followed by briefly cooling on ice, immediately before gel loading. Proteins were separated by SDS-PAGE on 10% Bolt Bis-Tris gels (catalog number NW00105BOX, Thermo Fisher Scientific). For PNGase F digested samples, 12% NuPage Bis-Tris gels (NP0342BOX, Thermo Fisher Scientific) were used. After the gel electrophoresis, proteins were transferred to 0.45 µm PVDF membranes (catalog number IPVH00010, Sigma-Aldrich, St. Louis, MO, USA). Following the blocking of the membrane in 5% skimmed milk (catalog number SKI400, BioShop Canada Inc) for 1 h at room temperature, membranes were incubated in primary antibodies overnight at 4°C with gentle rocking. Next, membranes were washed three times in 1x Tris-buffered saline containing 0.1% Tween-20 (TBST) (catalog number TWN508, BioShop) and incubated with the corresponding HRP-conjugated secondary antibodies for 1 h at room temperature. After washing the membranes trice again in 1x TBST, they were incubated for 1 min with Western Lightning Pro enhanced chemiluminescent (ECL) reagent (catalog number NEL120001EA, Revvity Health Sciences Inc., Mississauga, ON, Canada). Finally, membranes were exposed to autoradiography film (catalog number CLMS810, MedStore, Toronto, ON, Canada) and developed using a film developer.

### Sample preparation for mass spectrometry

The protein concentration of brain extracts was adjusted to 4 µg/µL. To denature all proteins in the sample, including the PrP^Sc^, 20 µg (in 5 uL) of total proteins were transferred to Protein LoBind (PLB, catalog number: PRE050LR-N, Diamed, Mississauga, ON, Canada) tubes and diluted with 9M urea (catalog number UR001.1, Bioshop) in a 1:2 ratio (v/v), achieving a 6 M urea concentration in the sample, followed by gentle vortexing and 30-minute incubation at room temperature to ensure complete denaturation. The pH of the samples was checked and, if below pH 8, increased to this pH using 1 M Triethylammonium Bicarbonate (TEAB) (catalog number 1861436, Thermo Fisher Scientific). Next the denatured proteins were reduced with 6.5 mM TCEP (catalog number TCE101, Bioshop) during a 30-minute incubation at 37°C, then alkylated for another 30 minutes in the dark at room temperature in the presence of 15 mM iodoacetamide (catalog number 1861445, Thermo Fisher Scientific).

Next, the proteins were subjected to solvent precipitation to maximize recovery for proteomic analyses ^69^. To this end, samples were threefold diluted to prepare them for the precipitation. In parallel, silica beads of 9–13 μm mean particle diameter (catalog number 440345), which had been pre-cleaned and resuspended in acetonitrile (ACN) at a concentration of 70 µg/µL, were added at a 10:1 bead-to-protein ratio and the tubes were gently vortexed. Subsequently, 100% ACN was added to each sample at a 1:4 (v/v) ratio to achieve an 80% ACN concentration without pipette mixing, followed by gentle vortexing for 10 seconds. The precipitates were centrifuged for 5 minutes at 16,000 × g (RCF) and room temperature, before supernatants were carefully removed, using the tube hinge as a guide to avoid disturbing the pellet. To further remove non-protein contaminants, the pellets were washed three times with 80% ethanol, using at least twice the total precipitation volume for each wash, followed by 2-minute centrifugation at 16,000 × g (RCF) and room temperature.

After the final wash, the remaining supernatant was carefully removed, and the protein pellet resuspended by gently vortexing in 100 mM ABC with a 1:50 trypsin-to-protein ratio. To fully disrupt the pellet, the tubes were placed in a sonication bath for 2 minutes. To ensure complete digestion, the samples were then incubated at 37°C for 18 hours at 800 RPM in a thermomixer (Thermomixer Comfort). Following digestion, 1 µL of 10% formic acid (FA) (catalog number A117-50, Thermo Fisher Scientific) was added to each sample to adjust the pH to approximately pH 3. The beads and any insoluble debris were precipitated by 10-minute centrifugation at 16,000 × g (RCF) and room temperature, before transferring the peptide-enriched supernatants to a new tube. To further recover additional peptides from the silica beads, the centrifugation pellets were washed once with 2% ACN in 0.1% formic acid, and the resultant wash supernatants combined with the initial peptide-containing supernatants. The final peptide concentration was adjusted to 0.25 to 0.5 µg/µL in 1% acetonitrile with 0.1% formic acid.

### Mass spectrometry data acquisition

Sample injected were three biological replicates from three treatment cohorts, with each sample composed of tryptic digests of 200 ng of total brain extract proteins. All data were acquired using a Vanquish Neo UHPLC system (Thermo Fisher Scientific) coupled to an Orbitrap Astral mass spectrometer (Thermo Fisher Scientific) through an EASY-Spray source (Thermo Fisher Scientific). Trap-and-Elute injection was based on PepMap Neo Trap Cartridge (catalog number 174500, Thermo Fisher Scientific) to filter impurities from peptide mapping samples. During peptide separation, a 15 cm EASY-Spray HPLC Column (catalog number ES900, Thermo Fisher Scientific) was maintained at 50°C. The mobile phase A consisted of 0.1% formic acid in water, while mobile phase B consisted of 0.1% formic acid and 80% acetonitrile in water. The gradient was as follows: the mobile phase was initially held at 2% B, increased from 2% to 4% in 0.5 min at a flow rate of 0.7 µL/min, then further increased to 5% B from 0.5 to 0.6 min, increased to 7% B from 0.6 to 1 min, ramped to 22.5% B from 1 to 18 min, then increased to 35% B from 18 to 25.5 min, and further increased to 50% B from 25.5 to 27 min. The flow rate was set to 0.5 µL/min. The column was then washed with 99% B from 27 to 30 min at a flow rate of 0.7 µL/min. Data were acquired in data independent acquisition (DIA) mode with a normalized collision energy of 25% and a default charge state of +2. MS1 spectra were acquired in the embedded orbitrap mass analyzer at a resolving power of 240,000 every 0.6 s. MS2 spectra were acquired in the embedded astral analyzer, with precursor isolation windows of 2 Th spanning the range of 380−980 Th. The custom normalized AGC target was set as 500% for both MS1 and DIA. The MS1 mass range was the same as the MS/ MS precursor range.

### Processing of global proteomic data set

Protein identification and peptide peak integration made use of Proteome Discoverer (PD) software (Version 3.2, Thermo Fisher Scientific) with CHIMERYS. These analyses interrogated the *Mus musculus* database (TaxID=10090, release=407, supplemented with the entry for the major prion protein from bank vole (Accession number Q8VHV5). The database search was restricted to tryptic peptides, allowing up to two missed cleavages per peptide. The maximum fragment mass tolerance was set to 10 ppm. Carbamidomethylation at cysteine residues was specified as a fixed modification, while oxidation at methionine residues was included as a variable modification. The sample-to-sample normalization in PD was based on the ‘Total Peptide Amount’ computed for each sample. Prion peptide comparisons were conducted using Skyline (Version 24.1) ^70^, with tubulin alpha and tubulin beta used for sample normalization. The assignment of peptide transitions to BvPrP were manually reviewed to ensure accurate peak integration. Both the Proteome Discoverer and Skyline data were exported to Excel for further conditional formatting.

### Cluster and KEGG analyses

Hierarchical clustering was undertaken with Cluster (Version 3.0, using Clustering Library Version 1.59) ^71, 72^. Prior to clustering, data columns were generated that inform about steady-state protein level ratios, computed by dividing cumulative MS1 ion intensities assigned to a given protein in a specific brain extract to the average MS1 ion intensities for the same protein in brain extracts from naïve mice. Subsequently, these steady-state protein level ratio data for all proteins, i.e., the rows within the Excel dataset, were clustered using centroid linkage clustering methods based on a ‘City-block distance’ metric and the relationship across the brain samples, i.e., the columns within the Excel dataset were inferred by hierarchical clustering using a Spearman Rank Correlation metric. Finally, hierarchical cluster analysis results were visualized in Java TreeView (Version 1.21) open source software ^71^.

KEGG pathway analyses were undertaken with the DAVID Bioinformatics suite (release DAVID 2021, Version 2023q4) of functional annotation tools made available by the National Institutes of Health ^73, 74^. Briefly, to initiate the analyses, lists of official gene symbols of genes whose expression gave rise to top- or bottom-ranked proteins in the global proteomic dataset were uploaded into the DAVID Analysis Wizard along with the selection of *Mus musculus* as the biological sample source. Next, a KEGG_PATHWAY analysis was undertaken from within the Annotation Summary Results page.

### Assignment of proteins increased in their steady-state levels in prion disease to mouse brain cell types

The cell type assignment made use of previously reported mouse brain cell type annotations, which were originally obtained through single-cell transcriptomic analyses ^40^. To interrogate these data with genes coding for proteins whose steady-state levels were most increased in RML-infected mice, which had reached the humane prion disease end-stage, we submitted the shortlist of genes to the online Transcriptomics Explorer algorithm (https://portal.brain-map.org/atlases-and-data/rnaseq). Separately, the ggplot package within RStudio was used to depict the non-neuronal portion of the brain cell type dendogram, which this analysis revealed to comprise the genes whose levels were most profoundly increased in RML prion disease.

### Statistical analyses

Western blot signal intensities from three biological replicates were quantified by densitometric analysis using ImageJ software and corresponding statistics were performed using Microsoft Excel. First, cohorts were subject to an F-test to assess the variance of the two compared groups. A Welch’s t-test or pooled t-test was subsequently chosen to assess the statistical significance of the differences between the averages of cohorts that were compared. Results were considered significant if p < 0.05 or denoted as non-significant with “ns” if p >0.05. Asterisks were used to signify varying levels of significance: p < 0.05 (*), p < 0.01 (**), and p < 0.001 (***).

Survival, weight, and nesting score charts were assembled in R using RStudio (Version 4.4.3). For all analyses, Excel raw data were uploaded into RStudio using the ‘readxl’ package. For the Kaplan-Meier analyses, the Greenwood formula computed a variance estimate, which was used by the Kaplan-Meier estimator (kmfit) to compute the 95% confidence interval. Plots were then generated with basic plotting functions embedded in ‘ggplot2’ and ‘ggfortify’ packages.

For generating the weight and nesting score charts, the geom_smooth() function within ggplot2 was used to depict the linear trend and the confidence interval using the Locally Estimated Scatterplot Smoothing ‘loess’ method.

We assumed a hypergeometric distribution when determining if it is statistically significant that 27 out of 30 cell surface proteins, which had been shown to reside in proximity to PrP in mouse brains ^42^, were downregulated ≥ 33% (out of 455 other proteins sharing this characteristic) in a global proteome dataset of 4874 proteins.

## REFERENCES

1. Prusiner SB. Prions. Proc Natl Acad Sci U S A 1998; 95: 13363–13383. 1998/11/13. DOI: 10.1073/pnas.95.23.13363.

2. Minikel EV, Vallabh SM, Lek M, et al. Quantifying prion disease penetrance using large population control cohorts. Sci Transl Med 2016; 8: 322ra329. 2016/01/23. DOI: 10.1126/scitranslmed.aad5169.

3. Stahl N, Borchelt DR, Hsiao K, et al. Scrapie prion protein contains a phosphatidylinositol glycolipid. Cell 1987; 51: 229–240. 1987/10/23.

4. Minikel EV, Zhao HT, Le J, et al. Prion protein lowering is a disease-modifying therapy across prion disease stages, strains and endpoints. Nucleic Acids Res 2020; 48: 10615–10631. 2020/08/11. DOI: 10.1093/nar/gkaa616.

5. Neumann EN, Bertozzi TM, Wu E, et al. Brainwide silencing of prion protein by AAV-mediated delivery of an engineered compact epigenetic editor. Science 2024; 384: ado7082. DOI: doi:10.1126/science.ado7082.

6. An M, Davis JR, Levy JM, et al. In vivo base editing extends lifespan of a humanized mouse model of prion disease. Nat Med 2025; 31: 1319–1328. 2025/01/15. DOI: 10.1038/s41591-024-03466-w.

7. Chou S-W, Mortberg MA, Marlen K, et al. Zinc Finger Repressors mediate widespread PRNP lowering in the nonhuman primate brain and profoundly extend survival in prion disease mice. bioRxiv 2025: 2025.2003.2005.636713. DOI: 10.1101/2025.03.05.636713.

8. Mead S, Whitfield J, Poulter M, et al. A novel protective prion protein variant that colocalizes with kuru exposure. N Engl J Med 2009; 361: 2056–2065. 2009/11/20. DOI: 10.1056/NEJMoa0809716.

9. Asante EA, Smidak M, Grimshaw A, et al. A naturally occurring variant of the human prion protein completely prevents prion disease. Nature 2015; 522: 478–481. 2015/06/11. DOI: 10.1038/nature14510.

10. Biffi A. Gene therapy for lysosomal storage disorders: a good start. Hum Mol Genet 2015; 25: R65–R75. DOI: 10.1093/hmg/ddv457.

11. Chesebro B, Trifilo M, Race R, et al. Anchorless prion protein results in infectious amyloid disease without clinical scrapie. Science 2005; 308: 1435–1439. 2005/06/04. DOI: 10.1126/science.1110837.

12. Stöhr J, Watts JC, Legname G, et al. Spontaneous generation of anchorless prions in transgenic mice. Proc Natl Acad Sci U S A 2011; 108: 21223–21228. 2011/12/14. DOI: 10.1073/pnas.1117827108.

13. Watts JC, Giles K, Bourkas ME, et al. Towards authentic transgenic mouse models of heritable PrP prion diseases. Acta Neuropathol 2016; 132: 593–610. 2016/06/29. DOI: 10.1007/s00401-016-1585-6.

14. Kocisko DA, Come JH, Priola SA, et al. Cell-free formation of protease-resistant prion protein. Nature 1994; 370: 471–474. 1994/08/11. DOI: 10.1038/370471a0.

15. Katorcha E, Makarava N, Savtchenko R, et al. Sialylation of prion protein controls the rate of prion amplification, the cross-species barrier, the ratio of PrPSc glycoform and prion infectivity. PLoS Pathog 2014; 10: e1004366. DOI: 10.1371/journal.ppat.1004366.

16. Hoyt F, Standke HG, Artikis E, et al. Cryo-EM structure of anchorless RML prion reveals variations in shared motifs between distinct strains. Nat Commun 2022; 13: 4005. DOI: 10.1038/s41467-022-30458-6.

17. Campana V, Caputo A, Sarnataro D, et al. Characterization of the properties and trafficking of an anchorless form of the prion protein. J Biol Chem 2007; 282: 22747–22756. 2007/06/09. DOI: 10.1074/jbc.M701468200.

18. Sangeetham SB, Engelke AD, Fodor E, et al. The G127V variant of the prion protein interferes with dimer formation in vitro but not in cellulo. Sci Rep 2021; 11: 3116. 2021/02/06. DOI: 10.1038/s41598-021-82647-w.

19. Huang JJ, Li XN, Liu WL, et al. Neutralizing Mutations Significantly Inhibit Amyloid Formation by Human Prion Protein and Decrease Its Cytotoxicity. J Mol Biol 2020; 432: 828–844. 2019/12/11. DOI: 10.1016/j.jmb.2019.11.020.

20. Hosszu LLP, Conners R, Sangar D, et al. Structural effects of the highly protective V127 polymorphism on human prion protein. Commun Biol 2020; 3: 402. 2020/07/31. DOI: 10.1038/s42003-020-01126-6.

21. Zheng Z, Zhang M, Wang Y, et al. Structural basis for the complete resistance of the human prion protein mutant G127V to prion disease. Sci Rep 2018; 8: 13211. 2018/09/06. DOI: 10.1038/s41598-018-31394-6.

22. Riek R, Hornemann S, Wider G, et al. NMR structure of the mouse prion protein domain PrP(121-231). Nature 1996; 382: 180–182. 1996/07/11. DOI: 10.1038/382180a0.

23. Donne DG, Viles JH, Groth D, et al. Structure of the recombinant full-length hamster prion protein PrP(29-231): the N terminus is highly flexible. Proc Natl Acad Sci U S A 1997; 94: 13452–13457. 1998/02/12. DOI: 10.1073/pnas.94.25.13452.

24. Zahn R, Liu A, Luhrs T, et al. NMR solution structure of the human prion protein. Proc Natl Acad Sci U S A 2000; 97: 145–150. 2000/01/05.

25. Gross DA, Tedesco N, Leborgne C, et al. Overcoming the Challenges Imposed by Humoral Immunity to AAV Vectors to Achieve Safe and Efficient Gene Transfer in Seropositive Patients. Front Immunol 2022; 13: 857276. 2022/04/26. DOI: 10.3389/fimmu.2022.857276.

26. Krieg AM, Yi AK, Matson S, et al. CpG motifs in bacterial DNA trigger direct B-cell activation. Nature 1995; 374: 546–549. 1995/04/06. DOI: 10.1038/374546a0.

27. Surabhi M, Matthew ECB, Lech K, et al. Convergent generation of atypical prions in knock-in mouse models of genetic prion disease. bioRxiv 2023: 2023.2009.2026.559572. DOI: 10.1101/2023.09.26.559572.

28. Watts JC, Giles K, Stohr J, et al. Spontaneous generation of rapidly transmissible prions in transgenic mice expressing wild-type bank vole prion protein. Proc Natl Acad Sci U S A 2012; 109: 3498–3503. 2012/02/15. DOI: 10.1073/pnas.1121556109.

29. Nonno R, Di Bari MA, Cardone F, et al. Efficient Transmission and Characterization of Creutzfeldt–Jakob Disease Strains in Bank Voles. PLoS Pathog 2006; 2: e12. DOI: 10.1371/journal.ppat.0020012.

30. Watts JC, Giles K, Patel S, et al. Evidence that bank vole PrP is a universal acceptor for prions. PLoS Pathog 2014; 10: e1003990. 2014/04/05. DOI: 10.1371/journal.ppat.1003990.

31. Fu H, Muenzer J, Samulski RJ, et al. Self-complementary adeno-associated virus serotype 2 vector: global distribution and broad dispersion of AAV-mediated transgene expression in mouse brain. Mol Ther 2003; 8: 911–917. 2003/12/11. DOI: 10.1016/j.ymthe.2003.08.021.

32. McCarty DM, Fu H, Monahan PE, et al. Adeno-associated virus terminal repeat (TR) mutant generates self-complementary vectors to overcome the rate-limiting step to transduction in vivo. Gene Ther 2003; 10: 2112–2118. 2003/11/20. DOI: 10.1038/sj.gt.3302134.

33. Gray SJ, Foti SB, Schwartz JW, et al. Optimizing promoters for recombinant adeno-associated virus-mediated gene expression in the peripheral and central nervous system using self-complementary vectors. Hum Gene Ther 2011; 22: 1143–1153. 2011/04/12. DOI: 10.1089/hum.2010.245.

34. Florea M, Nicolaou F, Pacouret S, et al. High-efficiency purification of divergent AAV serotypes using AAVX affinity chromatography. Mol Ther 2023; 28: 146–159. DOI: 10.1016/j.omtm.2022.12.009.

35. Nonnenmacher M, Wang W, Child MA, et al. Rapid evolution of blood-brain-barrier-penetrating AAV capsids by RNA-driven biopanning. Mol Ther 2021; 20: 366–378. 20201223. DOI: 10.1016/j.omtm.2020.12.006.

36. Zhao W, Eid S, Sackmann C, et al. Transient receptor potential vanilloid channel 2 contributes to multi-modal endoplasmic reticulum and perinuclear space dilations that can also be observed in prion-infected mice. bioRxiv 2024: 2024.2012.2018.629254. DOI: 10.1101/2024.12.18.629254.

37. Verkuyl C, Belotserkovsky A, Zerbes T, et al. Toward an all-in-one recombinant adeno-associated virus vector for functionally ablating the prion gene using CRISPR-Cas technology. PLoS One 2025; 20: e0336578. DOI: 10.1371/journal.pone.0336578.

38. Moreno JA, Radford H, Peretti D, et al. Sustained translational repression by eIF2alpha-P mediates prion neurodegeneration. Nature 2012; 485: 507–511. Research Support, Non- U.S. Gov’t 2012/05/25. DOI: 10.1038/nature11058.

39. Zhou S, Shi D, Liu X, et al. Protective V127 prion variant prevents prion disease by interrupting the formation of dimer and fibril from molecular dynamics simulations. Sci Rep 2016; 6: 21804. 2016/02/26. DOI: 10.1038/srep21804.

40. Yao Z, van Velthoven CTJ, Nguyen TN, et al. A taxonomy of transcriptomic cell types across the isocortex and hippocampal formation. Cell 2021; 184: 3222–3241.e3226. DOI: 10.1016/j.cell.2021.04.021.

41. Wang L, Cui C-Y, Lee CT, et al. Spatial transcriptomics of the aging mouse brain reveals origins of inflammation in the white matter. Nat Commun 2025; 16: 3231. DOI: 10.1038/s41467-025-58466-2.

42. Williams D, Mehrabian M, Arshad H, et al. The cellular prion protein interacts with and promotes the activity of Na,K-ATPases. PLoS One 2021; 16: e0258682. DOI: 10.1371/journal.pone.0258682.

43. Zulianello L, Kaneko K, Scott M, et al. Dominant-negative inhibition of prion formation diminished by deletion mutagenesis of the prion protein. J Virol 2000; 74: 4351–4360. 2001/02/07. DOI: 10.1128/jvi.74.9.4351-4360.2000.

44. Kaneko K, Zulianello L, Scott M, et al. Evidence for protein X binding to a discontinuous epitope on the cellular prion protein during scrapie prion propagation. Proc Natl Acad Sci U S A 1997; 94: 10069–10074. 1997/09/18.

45. Perrier V, Kaneko K, Safar J, et al. Dominant-negative inhibition of prion replication in transgenic mice. Proc Natl Acad Sci U S A 2002; 99: 13079–13084. 2002/09/25. DOI: 10.1073/pnas.182425299.

46. Kishida H, Sakasegawa Y, Watanabe K, et al. Non-glycosylphosphatidylinositol (GPI)-anchored recombinant prion protein with dominant-negative mutation inhibits PrPSc replication in vitro. Amyloid 2004; 11: 14–20. 2004/06/10. DOI: 10.1080/13506120410001689634.

47. Furuya K, Kawahara N, Yamakawa Y, et al. Intracerebroventricular delivery of dominant negative prion protein in a mouse model of iatrogenic Creutzfeldt-Jakob disease after dura graft transplantation. Neurosci Lett 2006; 402: 222–226. DOI: 10.1016/j.neulet.2006.03.062.

48. Toupet K, Compan V, Crozet C, et al. Effective Gene Therapy in a Mouse Model of Prion Diseases. PLoS One 2008; 3: e2773. DOI: 10.1371/journal.pone.0002773.

49. Arshad H, Patel Z, Amano G, et al. A single protective polymorphism in the prion protein blocks cross-species prion replication in cultured cells. J Neurochem 2023; 165: 230–245. 2022/12/14. DOI: 10.1111/jnc.15739.

50. Gatdula JRP, Orbe IC, Tolton SG, et al. Leveraging the dominant-negative effect of the kuru-protective G127V prion protein variant as a novel therapeutic strategy. bioRxiv 2026: 2026.2002.2017.703887. DOI: 10.64898/2026.02.17.703887.

51. Gatdula JRP, Mercer RCC, Alepuz Guillen JA, et al. Membrane-anchored PrPSc is the trigger for prion synaptotoxicity. PLOS Pathog 2026; 22: e1013911. DOI: 10.1371/journal.ppat.1013911.

52. Song F, Kovac V, Mohammadi B, et al. Cleavage site-directed antibodies reveal the prion protein in humans is shed by ADAM10 at Y226 and associates with misfolded protein deposits in neurodegenerative diseases. Acta Neuropathol 2024; 148: 2. 2024/07/09. DOI: 10.1007/s00401-024-02763-5.

53. Song F, Kovac V, Mohammadi B, et al. Proteolytic shedding of the prion protein: Uncovering “new” biological implications of a conserved cleavage event. Neural Regen Res 2025 2025/07/10. DOI: 10.4103/nrr.Nrr-d-25-00013.

54. Vanni I, Iacobone F, D’Agostino C, et al. An optimized Western blot assay provides a comprehensive assessment of the physiological endoproteolytic processing of the prion protein. J Biol Chem 2023; 299: 102823. DOI: 10.1016/j.jbc.2022.102823.

55. Kordek R, Liberski PP, Yanagihara R, et al. Molecular analysis of prion protein (PrP) and glial fibrillary acidic protein (GFAP) transcripts in experimental Creutzfeldt-Jakob disease in mice. Acta Neurobiol Exp 1997; 57: 85–90. 1997/01/01.

56. Lauren J, Gimbel DA, Nygaard HB, et al. Cellular prion protein mediates impairment of synaptic plasticity by amyloid-beta oligomers. Nature 2009; 457: 1128–1132. 2009/02/27. DOI: 10.1038/nature07761.

57. Barry AE, Klyubin I, Mc Donald JM, et al. Alzheimer’s disease brain-derived amyloid-beta-mediated inhibition of LTP in vivo is prevented by immunotargeting cellular prion protein. J Neurosci 2011; 31: 7259–7263. 2011/05/20. DOI: 10.1523/JNEUROSCI.6500-10.2011.

58. Freir DB, Nicoll AJ, Klyubin I, et al. Interaction between prion protein and toxic amyloid beta assemblies can be therapeutically targeted at multiple sites. Nat Commun 2011; 2: 336. 2011/06/10. DOI: 10.1038/ncomms1341.

59. Rushworth JV, Griffiths HH, Watt NT, et al. Prion protein-mediated toxicity of amyloid-beta oligomers requires lipid rafts and the transmembrane LRP1. J Biol Chem 2013; 288: 8935–8951. Research Support, Non-U.S. Gov’t 2013/02/07. DOI: 10.1074/jbc.M112.400358.

60. Ganzinger KA, Narayan P, Qamar SS, et al. Single-molecule imaging reveals that small amyloid-beta1-42 oligomers interact with the cellular prion protein (PrP(C)). Chembiochem 2014; 15: 2515–2521. 2014/10/09. DOI: 10.1002/cbic.201402377.

61. Purro SA, Nicoll AJ and Collinge J. Prion Protein as a Toxic Acceptor of Amyloid-beta Oligomers. Biol Psychiatry 2018; 83: 358–368. 2018/01/15. DOI: 10.1016/j.biopsych.2017.11.020.

62. Gunther EC, Smith LM, Kostylev MA, et al. Rescue of Transgenic Alzheimer’s Pathophysiology by Polymeric Cellular Prion Protein Antagonists. Cell Rep 2019; 26: 1368. 2019/01/31. DOI: 10.1016/j.celrep.2019.01.064.

63. Gimbel DA, Nygaard HB, Coffey EE, et al. Memory impairment in transgenic Alzheimer mice requires cellular prion protein. J Neurosci 2010; 30: 6367–6374. 2010/05/07. DOI: 30/18/6367 [pii] 10.1523/JNEUROSCI.0395-10.2010.

64. Horn N, Tomlinson A, Jakobcova T, et al. Exploring the protective roles of prion protein polymorphisms in novel gene targeted mouse models. 2025 International Prion Meeting 2025; Abstract.

65. Hollidge BS, Carroll HB, Qian R, et al. Kinetics and durability of transgene expression after intrastriatal injection of AAV9 vectors. Front Neurol 2022; 13: 1051559. 2022/12/02. DOI: 10.3389/fneur.2022.1051559.

66. Williamson RA, Peretz D, Pinilla C, et al. Mapping the prion protein using recombinant antibodies. J Virol 1998; 72: 9413–9418. 1998/10/10.

67. Mehra S, Bourkas ME, Kaczmarczyk L, et al. Convergent generation of atypical prions in knockin mouse models of genetic prion disease. J Clin Invest 2024; 134 2024/08/01. DOI: 10.1172/jci176344.

68. Kawamoto T and Kawamoto K. Preparation of Thin Frozen Sections from Nonfixed and Undecalcified Hard Tissues Using Kawamoto’s Film Method (2020). In: Hilton MJ (ed) Skeletal Development and Repair: Meth Protocols. New York, NY: Springer US, 2021, pp.259–281.

69. Johnston HE, Yadav K, Kirkpatrick JM, et al. Solvent Precipitation SP3 (SP4) Enhances Recovery for Proteomics Sample Preparation without Magnetic Beads. Anal Chem 2022; 94: 10320–10328. 2022/07/19. DOI: 10.1021/acs.analchem.1c04200.

70. Pino LK, Searle BC, Bollinger JG, et al. The Skyline ecosystem: Informatics for quantitative mass spectrometry proteomics. Mass Spectrom Rev 2020; 39: 229–244. 2017/07/12. DOI: 10.1002/mas.21540.

71. Eisen MB, Spellman PT, Brown PO, et al. Cluster analysis and display of genome-wide expression patterns. Proc Natl Acad Sci USA 1998; 95: 14863–14868. 1998/12/09.

72. de Hoon MJ, Imoto S, Nolan J, et al. Open source clustering software. Bioinformatics 2004; 20: 1453–1454. 2004/02/12. DOI: 10.1093/bioinformatics/bth078.

73. Huang da W, Sherman BT and Lempicki RA. Systematic and integrative analysis of large gene lists using DAVID bioinformatics resources. Nat Protoc 2009; 4: 44–57. 2009/01/10. DOI: 10.1038/nprot.2008.211.

74. Sherman BT, Hao M, Qiu J, et al. DAVID: a web server for functional enrichment analysis and functional annotation of gene lists (2021 update). Nucleic Acids Res 2022; 50: W216–w221. 2022/03/25. DOI: 10.1093/nar/gkac194.

